# Transcriptome analysis of *Plasmodium falciparum* isolates from Benin reveals specific gene expression associated with cerebral malaria

**DOI:** 10.1101/2021.11.08.467248

**Authors:** E. Guillochon, J. Fraering, V. Joste, C. Kamaliddin, B. Vianou, L. Houzé, L.G Baudrin, J.F. Faucher, A. Aubouy, S. Houzé, M. Cot, N. Argy, O. Taboureau, G.I. Bertin, NeuroCM group

## Abstract

The host and parasitic factors leading to cerebral malaria (CM) are not yet fully elucidated and CM *Plasmodium falciparum* isolates transcriptome profile remains largely unknown. Based on RNA-seq data from 15 CM and 15 uncomplicated malaria (UM) children from Benin, we identified an increased ring stage signature in CM parasites. Reduced circulating time may result from a higher adherence ability of CM isolates and consistent with this hypothesis, we measured an overexpression of *var* genes in CM. *var* genes domains expression was more restricted in CM isolates compared to UM, reflecting the specific binding to receptors in host brain endothelium capillaries. However, ICAM-1 binding motif was found expressed in both CM and UM, questioning its role in *Pf*EMP1 adhesion to ICAM-1 receptor. UM isolates increased circulation time may also be modulated by a more efficient immune response against infected erythrocytes surface proteins, which we could not demonstrate on our cohort. Identification of deregulated genes involved in adhesion, excluding variant surface antigens, also supports the hypothesis of an increased CM adhesion capacity. Finally, numerous upregulated genes involved in entry into host pathway were found, reflecting a greater erythrocytes invasion capacity of CM parasites.

## Introduction

Malaria is a vector-borne disease that affected 229 million people in 2019 (1). *Plasmodium falciparum* species is responsible for most cases and deaths in sub-Saharan Africa. Cerebral malaria (CM) is the most severe form of *Plasmodium falciparum* infections and mainly affects children under five years old, who account for nearly 67% of malaria deaths (1). CM pathophysiology is related to *P. falciparum* ability to adhere efficiently to specific receptors in host brain capillaries (2,3), during asexual intraerythrocytic development cycle (IDC). Throughout this cycle, the parasite evolves from ring, young to late trophozoite, schizont and finally merozoite form. Erythrocytes infected with *P. falciparum* mature forms (*i.e*. trophozoites and schizonts) can adhere to the host endothelium, through variant surface antigens (VSA) exported to the erythrocyte membrane (4). Recent studies showed that infected erythrocytes (iE) circulation time is decreased in severe malaria, which reflects a greater adherence of mature parasitic forms to endothelial vessels (5–8). Higher adherence capacity confers a real advantage to the parasite by protecting iE from splenic clearance thus promoting increased parasitaemia (8). This phenomenon is modulated by both the host’s immune system (9) and iE intrinsic adherence capacity, mainly mediated by *Plasmodium falciparum* Erythrocyte Membrane Protein 1 (*Pf*EMP1) surface antigen.

*Pf*EMP1 is the major VSA and is encoded by the multigenic *var* genes family (10). *Pf*EMP1 contain two types of hypervariable extracellular domains: cysteine-rich interdomain regions (CIDR) and Duffy-binding-like (DBL), classified into subdomains based on amino acid sequence. Some specific subdomains enchainment are defined as domain cassette (DC) (11), among which DC8 and DC13 are associated with CM (12,13). The ability of these DCs to bind to Endothelial Protein C Receptor (EPCR) *via* the CIDRα1 domain is a major virulence factor of malaria severe forms (2). Moreover, *Pf*EMP1 with dual-binding activities were defined by the combination of a CIDRα1 domain binding EPCR followed by a DBLβ1/3 domain presenting a dedicated amino acid binding motif, and were associated with CM pathophysiology (14,15).

The present work is part of the NeuroCM study, investigating host and parasite factors leading to neuroinflammation in CM. For this purpose, children presenting with CM and uncomplicated malaria (UM) were recruited in Benin in 2018, during the malaria transmission season (16). A first comparative study (17) highlighted that CM isolates display a greater adherence to Hbec-5i and CHO-ICAM-1 cell lines than UM. The overexpression of EPCR binding CIDRα1 domains was confirmed in CM isolates by RT-qPCR (17). However, DBLβ1/3 domains containing ICAM-1 binding motif were found in both groups and were not overexpressed in CM isolates compared to UM, questioning its role in CM pathophysiology.

While the *var* genes domains expression associated with CM has been extensively studied (13,17,18), the others parasitic factors causing CM are not yet fully understood. Previous studies focusing on severe malaria (SM) parasite transcriptome have shown that parasites had a specific gene expression compared to those infecting patients presenting UM (5,6). A more metabolically quiescent phenotype associated with SM has been demonstrated, resulting from deregulated transcripts in the glycolytic, folate and pyrimidine biosynthesis pathways (5). In addition, deregulated genes implicated in the *var* genes expression or *Pf*EMP1 presentation to iE membrane were identified (5,6).

More in-depth transcriptomic analyzes could unravel specific parasite gene expression associated with malaria severity. In this context, we aimed to characterize *P. falciparum* gene expression associated with CM in Beninese children. To this end, RNA-sequencing of 15 CM and 15 UM isolates was performed. This study showed that iE circulation time is shorter in CM isolates, due to an increased adherence ability of CM parasites. Based on semi-conserved sequences, we were able to measure a *var* genes overexpression in CM isolates. Finally, we focused on CM gene expression specificities through a differential expression approach.

## Results

### Clinical and biological characteristics comparison highlights a difference in parasitaemia and age between CM and UM children

15 CM and 15 UM children were selected. They were chosen based on the quantification and quality of parasite RNA and RNA-seq mapping statistics (supplemental data 1 and 2). Clinical and biological characteristics of these patients are described in Table 1. Patients with CM had a higher parasitaemia (p<0.001, Mann-Whitney U-test) and UM children were older (p=0.004, Mann-Whitney U-test). Total antibodies titers evaluated by indirect immunofluorescence (p=0.57, χ^2^ test), as well as total IgG quantification measured by flow cytometry (42,575 [23,046-87,243] *vs* 28,721 [19,148-55,502] RFU, p=0.12, Mann-Whitney U-test) in patient plasma and against 3D7 schizonts iE were not different between CM and UM (supplemental data 3).

**Table 1:** Clinical and biological features of the sequenced and additional groups. Quantitative variables are presented by their median [10^th^ - 90^th^ percentiles]. Statistical measurements were assessed with Mann-Whitney U-test. Qualitative variables are presented by proportions and statistical differences were measured with the χ^2^ test.

|  | Sequenced group |  |  | Additional group |  |  |
| --- | --- | --- | --- | --- | --- | --- |
|  | CM (n=15) | UM (n=15) | p-value | CM (n=15) | UM (n=15) | p-value |
| **Age** | 42.4 | 55 | 0.004 | 40.8 | 50.4 | NS |
| **(months)** | [30.2-55.5] | [44.5-70.1] |  | [32.2-63.36] | [24.2-64.321] |  |
| **Gender:** |  |  |  |  |  |  |
| **female no /total no** | 6/15 (40%) | 4/15 (27%) | NS | 10/15 (67%) | 7/15 (47%) | NS |
| **(% female)** |  |  |  |  |  |  |
| **Parasitaemia** | 459,000 | 252,000 | <0.001 | 63,000 | 64,527 | NS |
| **(parasites/μl)** | [281,600-1152,286] | [148,542-325,626] |  | [3,745-403,200] | [6,539-155,604] |  |
| **Hemoglobinemia (g/dl)** | 6.5 [3.78-9.18] | 9.8 [8.34-10.68] | <0.001 | 5.6 [3.24-8.46] | 8.6 [7.32-10.22] | <0.001 |
| **Blood glucose** | 0.14 [0.1-0.71] | 1.05 [0.82-1.36] | <0.001 | 0.9 [0.1-1.28] | 1 [0.83-1.38] | NS |
| **(g/liter)** |  |  |  |  |  |  |
| **Malarial retinopathy** | Yes: 5<br>No: 4<br>Not evaluated: 6 | NA | NA | Yes: 7<br>No: 5<br>Not evaluated: 3 | NA | NA |
|  | 0: 1 (6.7%) |  |  | 0: 0 (0%) |  |  |
| **Blantyre score** | 1: 5 (33.3%) | NA | NA | 1: 6 (40%) | NA | NA |
|  | 2: 9 (60%) |  |  | 2: 9 (60%) |  |  |
| **Mortality (%)** | 57% | NA | NA | 18% | NA | NA |

We designed an additional group of 15 CM and 15 UM from the same cohort to corroborate the major results observed on the sequenced group (Table 1). In this group, there were no significant difference in parasitaemia (p=0.39, Mann-Whitney U-test) or age (p=0.14, Mann-Whitney U-test) between the clinical phenotypes. Parasitaemia was significantly higher in the sequenced group compared to additional in both CM (p<0.001, Mann-Whitney U-test) and UM phenotypes (p<0.001, Mann-Whitney U-test). Blood glucose was lower in the sequenced group compared to the additional group for CM (p=0.02, Mann-Whitney U-test).

### CM children have an increased proportion of circulating ring stage parasites

The all-gene analysis was focused on parasite core genome (VSAs were excluded). Parasite developmental stages proportions were evaluated in each sample based on total gene expression measured by RNA-seq, using a previously published algorithm (5) (Figure 1A, supplemental data 4). The mean parasite’s age expressed in hours post infection (hpi) was statistically lower in CM cases, with a median age of 8 hpi [8-11.2] compared to 12.6 hpi [8.8-16.7] in UM (p<0.001, Mann-Whitney U-test). The difference was confirmed by the microscopic reading of thin blood smears (p<0.001, Mann-Whitney U-test) (supplemental data 4). In the additional group, parasites’ stage proportions were evaluated by microscopic reading (Figure 1A, supplemental data 4), confirming the difference highlighted in the sequenced group (p=0.05, Mann-Whitney U-test). Before correcting gene expression for life-cycle stage effect through published algorithm (5), the all-gene expression principal-component analysis (PCA) showed that both clinical presentation and parasite developmental stage affect the separation by the first component (Figure 1B). In fact, samples 03-022 and 03-113 exhibited a majority of ring stage parasites and grouped together with the CM samples. After the correction, CM and UM isolates were perfectly separated (Figure 1B).

**Figure 1:**
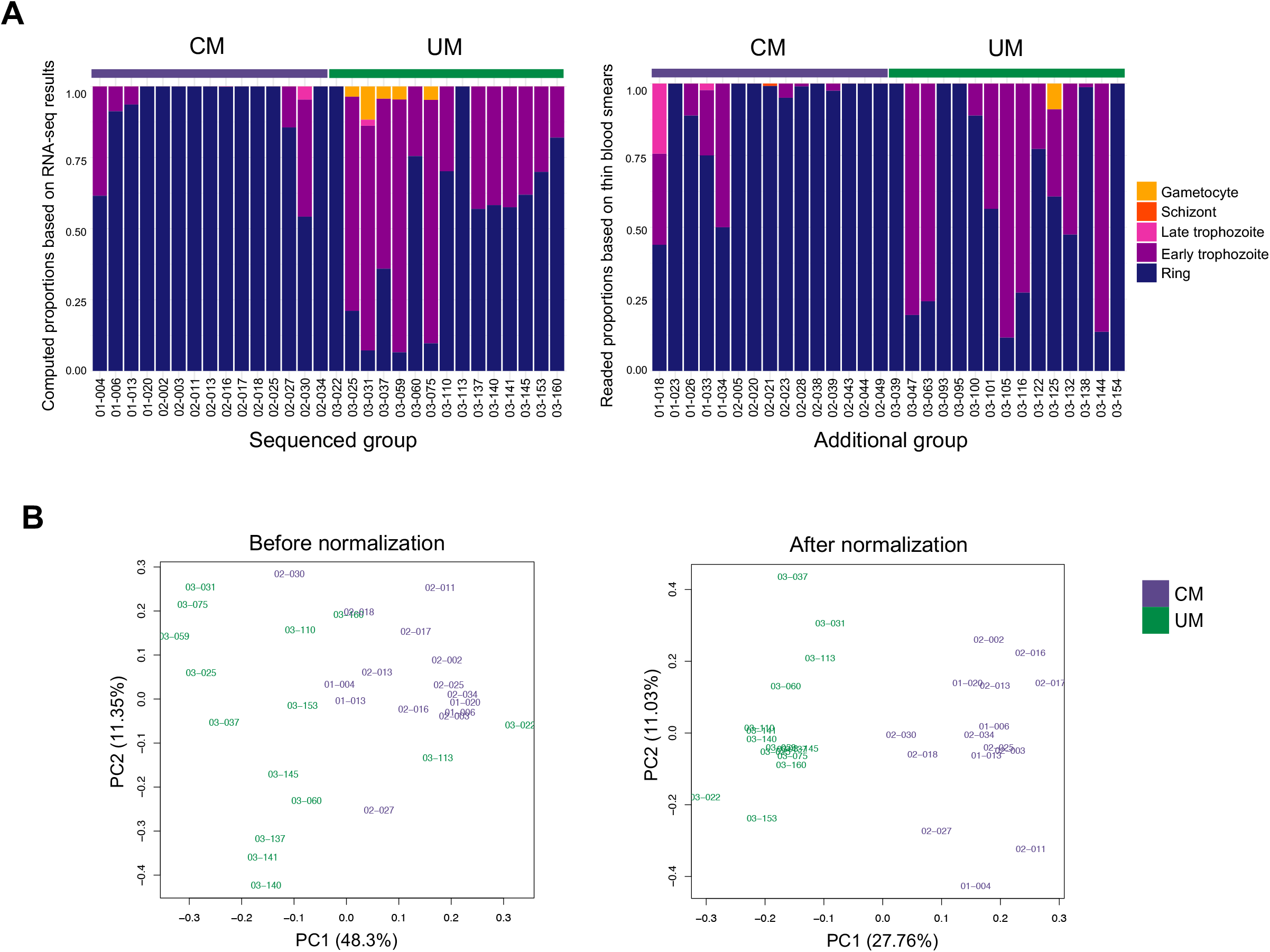
Parasites developmental stage proportions and PCA computed from all-gene expression. **A/** Proportion of ring, early trophozoite, late trophozoite, schizont and gametocyte stages parasites in each sample. In the sequenced group, stage proportions were evaluated based on gene expression measured by RNA-seq. In the additional group, they were determined by microscopic reading of thin blood smears. **B/** Principal Component Analysis based on the all-gene expression before normalization (left) and after normalization (right) for life cycle biases.

### Parasite gene expression is a signature of malaria severity

The differential expression analysis revealed 551 differentially expressed (DE) genes (supplemental table 1), of which 284 were upregulated and 267 were downregulated in CM compared to UM (absolute fold-change (FC)>1.5 and p≤0.05). The top 20 DE genes based on absolute FC values are displayed in Table 2. Further hierarchical clustering of the DE genes showed that CM and UM isolates were grouped together, reflecting a common gene expression pattern within each clinical group (Figure 2A).

**Figure 2:**
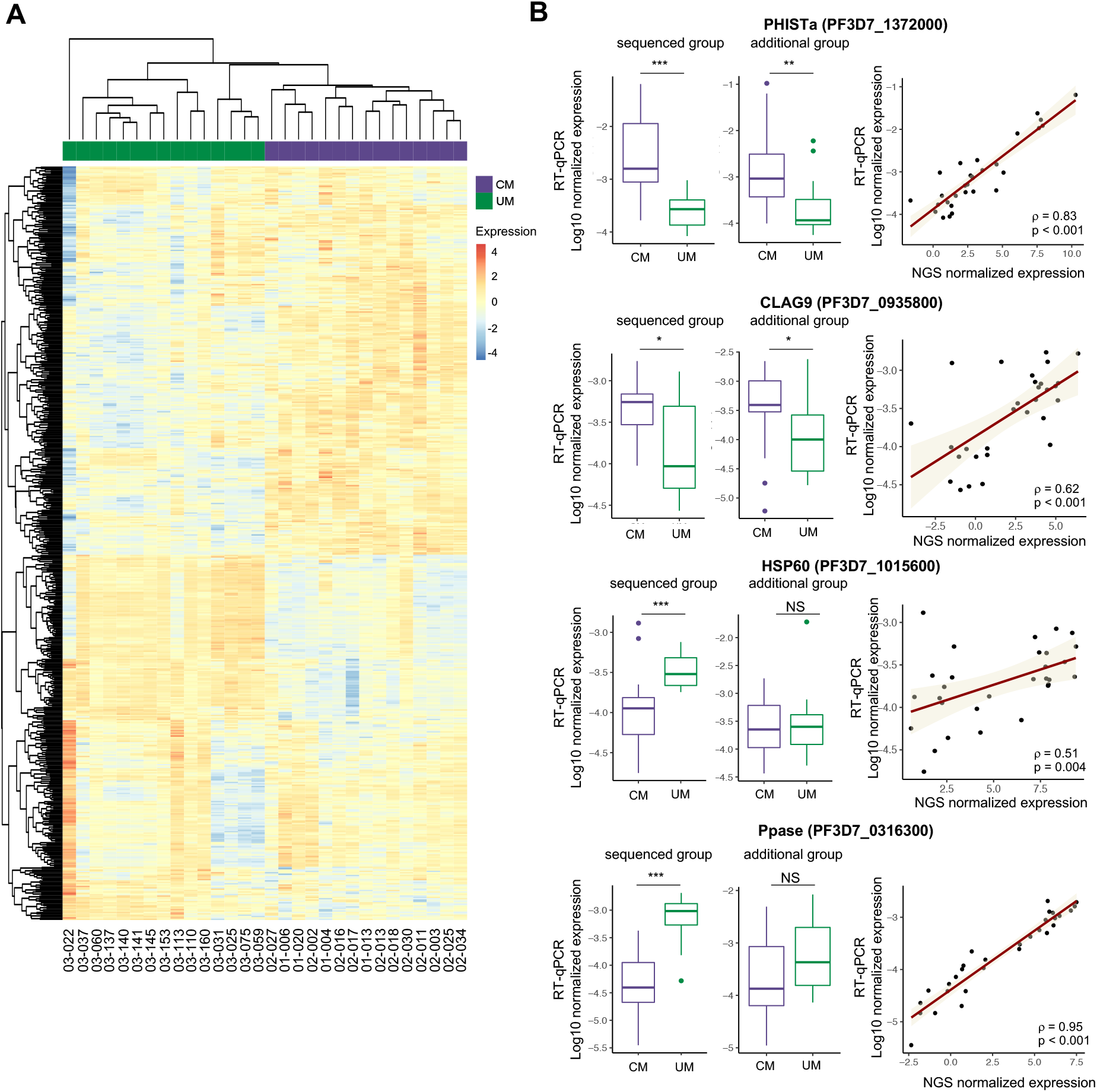
Differential expression analysis and RT-qPCR validations. **A/** Heatmap of differentially expressed genes. An upregulated expression is represented in red and the downregulation in blue. **B/** Distribution of normalized expression values measured by RT-qPCR in CM and UM is represented (boxplot). Results are displayed for both the sequenced and the additional group. The p-values were calculated with the Mann-Whitney U-test and are represented by: *** <0.001; ** <0.01; * <0.05. The correlation between the normalized expression values measured by RT-qPCR and RNA-seq was represented through the regression line and tested by the Spearman rank test (Spearman rank correlation coefficient ρ and p-value are shown).

**Table 2:**
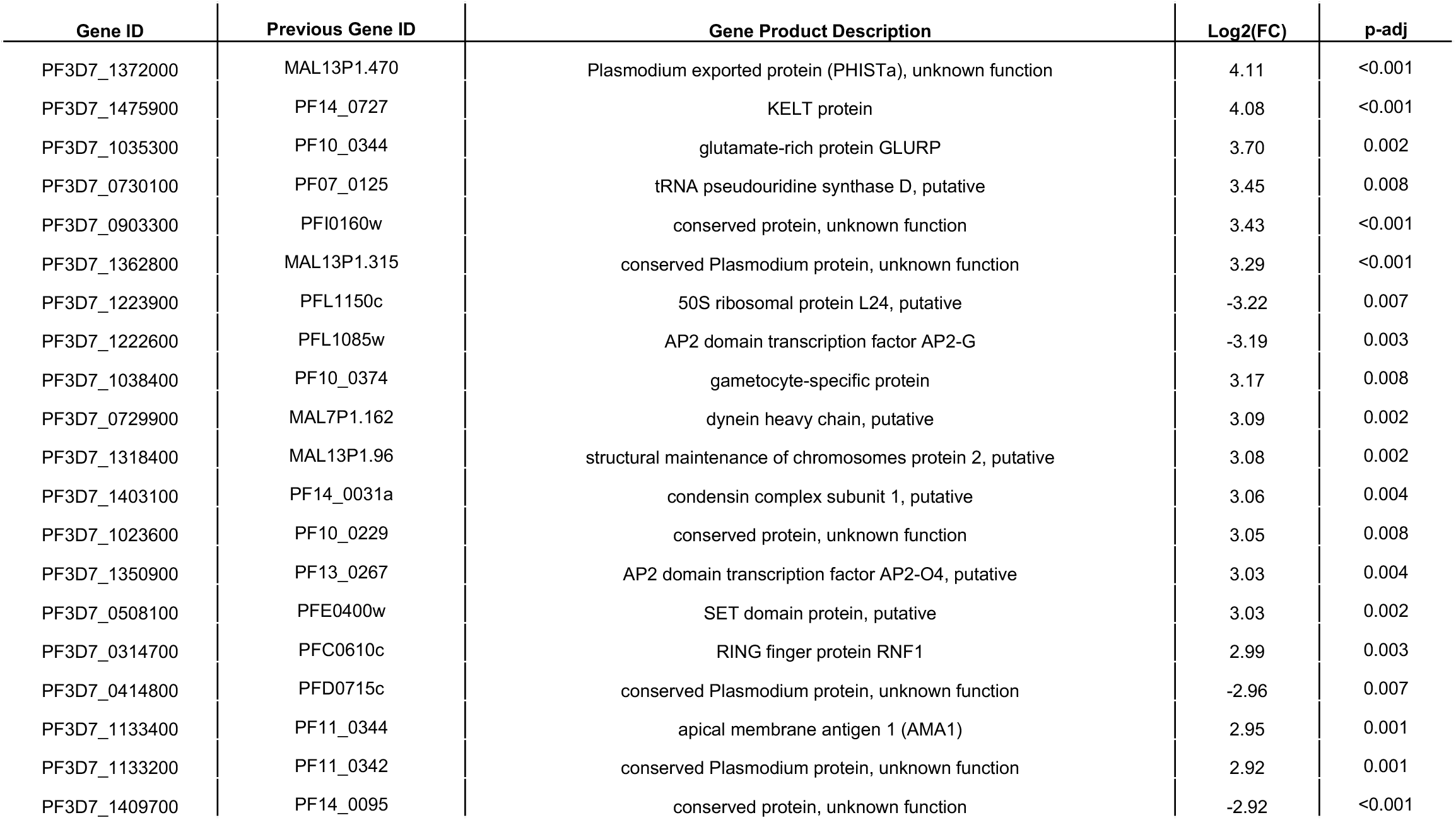
Top 20 deregulated genes based on absolute FC value. We have provided the current and the previous gene IDs, and the gene product description. A positive Log2-transform fold-change (Log2(FC)) value is associated with an upregulated expression in CM compared to UM. A negative value is associated with a downregulated expression in CM. Adjusted p-values (Benjamini-Hochberg) are given.

We chose eight DE genes to confirm differential gene expression measured by RNA-seq, four genes were upregulated in CM and four were downregulated (supplemental data 5). The corresponding proteins have variable functions and locations. Gene expression was corrected by both housekeeping gene expression and IDC stage proportions. We show in Figure 2B the results for two upregulated genes: PHISTa (PF3D7_1372000, FC=17.2, p-adj<0.001), CLAG9 (F3D7_0935800, FC=2.3, p-adj=0.04) and two downregulated genes: HSP60 (PF3D7_1015600, FC=-3.3, p-adj=0.002), Ppase (PF3D7_0316300, FC=-4.7, p-adj=0.02). For the upregulated genes, the difference was validated in the sequenced and additional group. For downregulated genes, the result was validated for the sequenced group but not for the additional group. Similar results were observed for the other 2 up-regulated and down-regulated genes (supplemental data 5). In the additional group, we did not measure a significant difference for the downregulated genes but for Ppase (p=0.19, Mann-Whitney U-test) and ARFGAP2 (p=0.2, Mann-Whitney U-test) we observed a tendency to downregulation. For all eight genes, RT-qPCR expression was strongly correlated with gene expression measured by RNA-seq, with all Spearman rank correlation coefficients ρ greater than 0.5 (all p<0.01, Spearman rank test).

### Gene set enrichment analysis (GSEA) of DE genes shows deregulation in pathways related to metabolism and entry into host

Gene Ontology (GO) and Kyoto Encyclopedia of Genes and Genomes (KEGG) pathways enrichment analysis of all the DE genes showed 13 deregulated GO pathways (biological process), with a minimum of 18% deregulated genes identified in each (p≤0.05, hypergeometric test) (figure 3A, supplemental data 6). We found three deregulated pathways that were part of DNA metabolic processes, two pathways were involved in protein metabolic processes and three pathways implicated in organelle organization. Entry into host pathway was also deregulated, including 12 up-regulated genes in CM and 2 down-regulated genes, of which apical membrane antigen 1 (AMA-1, PF3D7_1133400), Plasmepsin IX and X (PMIX, PF3D7_1430200 and PMX, PF3D7_0808200) and erythrocyte binding antigen-175 (eba-175, PF3D7_0731500) were upregulated. Concerning KEGG pathways results, we found three associated pathways, with at least 40% deregulated genes, including pyruvate metabolism and glycolysis/gluconeogenesis.

**Figure 3:**
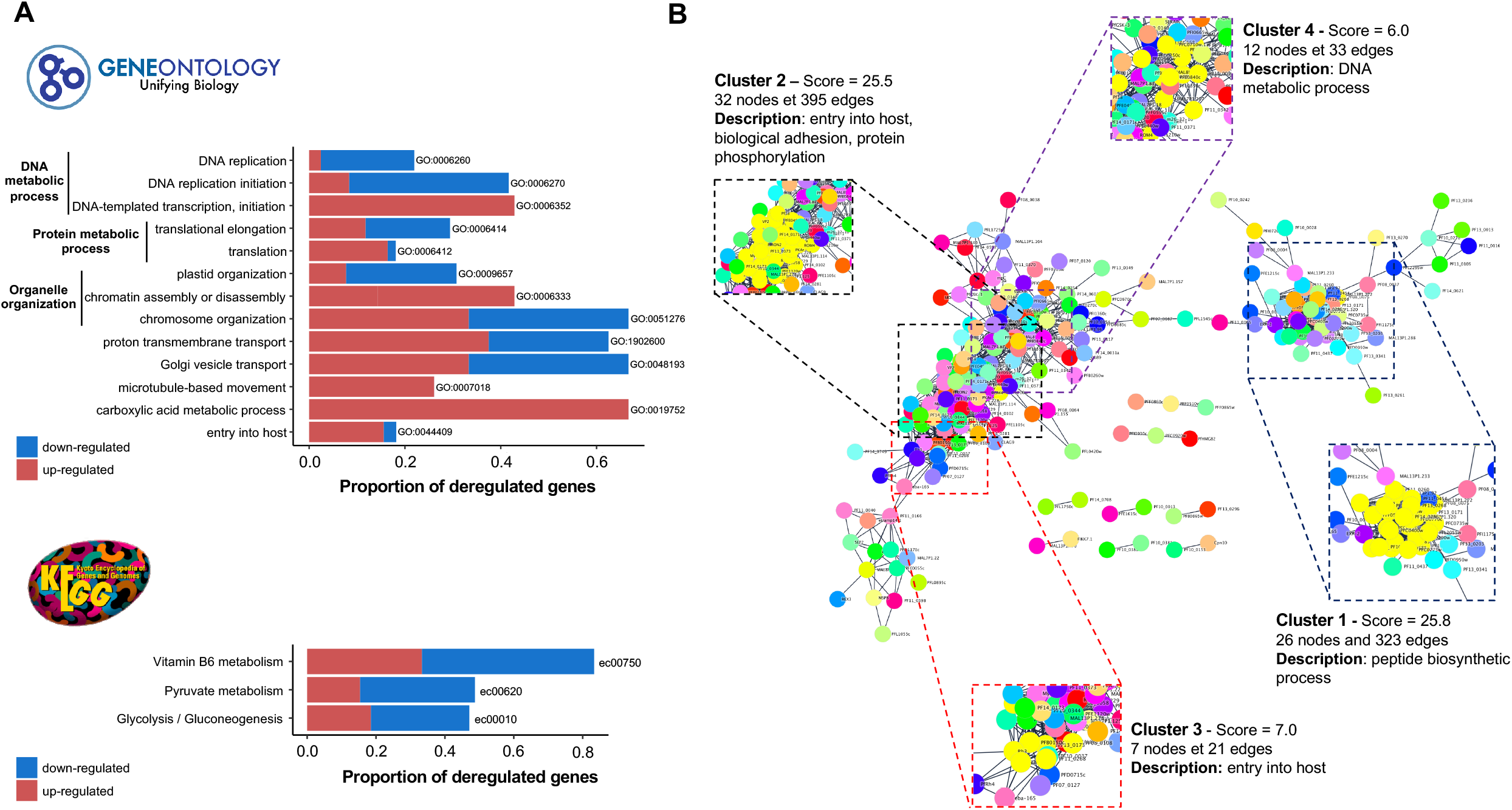
GSEA and PPI network of DE genes analysis. **A**/ GO and KEGG pathways enrichment analysis. Barplots show the proportion of deregulated genes in each pathway. Upregulated genes in CM are in red and downregulated in CM in blue. **B**/ String’s PPI network of DE genes. Cluster found with MCode and associated pathways are shown.

### Protein–protein interaction (PPI) network analysis reveals highly connected clusters involved in entry into host and cytoadhesion

We performed a PPI network analysis to demonstrate the functional relationships between DE genes. 546 DE genes were annotated in the STRING database (out of 551) and could therefore be analyzed. Results showed a very dense network with 212 nodes and 2,976 edges with a significant PPI enrichment (p<0.001), reflecting biologically connected proteins (Figure 3B). For a search of highly interconnected regions, MCode was used, and 4 clusters were found (Figure 3B, supplemental data 7). The cluster 1, containing 26 nodes and 323 edges, was composed exclusively of ribosomal proteins (all downregulated), reflecting the peptide biosynthesis process deregulation. In cluster 2, there were 32 nodes and 395 edges, and the main pathways represented in this cluster were entry into host (rhoptry-associated protein 2 (PF3D7_0501600), AMA-1 (PF3D7_1133400) and 6-cysteine protein (PF3D7_0508000), biological adhesion (CLAG proteins) and protein phosphorylation. In cluster 3, composed of 7 nodes and 21 edges, proteins were implicated in entry into host through erythrocyte binding antigen-181 (PF3D7_0102500) and PMX (PF3D7_0808200). Cluster 4 contained 12 nodes and 33 edges represented activities related to DNA metabolic process.

### Genes implicated in *var* genes expression regulation, *Pf*EMP1 trafficking and presentation to membrane are differentially expressed between CM and UM isolates

As genes involved in the *Pf*EMP1 expression process are not part of an annotated pathway, we manually identified them. In the top 20 DE genes (Table 2), we identified upregulated genes including PHISTa (PF3D7_1372000/MAL13P1.470), SET9, (PF3D7_0508100), SET10 (PF3D7_1221000) and ApiAP2 (PF3D7_1350900). In the rest of the set, we also identified upregulated genes as PHISTb (PF3D7_0424600), FIKK9.6 (PF3D7_0902500) and SEMP1 (PF3D7_0702400) and downregulated genes as EXP2 and EXP3 (PF3D7_1471100; PF3D7_1024800).

### *var* genes were assembled, and domains expression comparison shows a grouping of CM isolates

*var* genes from each isolate were assembled from RNA-seq reads based on a previously published pipeline (5). As a reminder, we have represented the schematic structure of *Pf*EMP1 in the Figure 4A. The majority (28/30) of the sequenced samples were analyzed with at least one *var* domain identified. A higher number of *var* assemblies was obtained in CM isolates with 96.0 [70.4-141.2] assemblies with at least one predicted *Pf*EMP1 domain compared to 41.0 [2.0-102] in UM samples (p=0.002, Mann-Whitney U-test) (Figure 4B). As we obtained more *P. falciparum* mapped reads in CM isolates compared to UM (21,589,730 [14,293,869-29,416,042] *vs* 15,826,946 [10,424,471-19,131,881], p=0.016, Mann-Whitney U-test), this could be the cause of the difference in the number of assemblies obtained. We sought to assess the quality of the assemblies in each patient by comparing reads aligned to the semi-conserved ATS domain (exon2) to the number of reads aligned to the exon 1 assemblies (figure 4C). The proportion of well reconstructed domains was higher in CM compared to UM (0.57 [0.48-0.7] *vs* 0.39 [0.04-0.64], p=0.007, Mann-Whitney U-test). After removal of the five isolates with an exon1 / ATS ratio less than 0.3 (03-031, 03-037, 03-059, 03-075 and 03-137), the median ratios were similar between CM and UM (0.57 [0.48-0.7] *vs* 0.50 [0.29-0.68], p=0.33, Mann-Whitney U-test). If we then count the number of contigs with at least one domain reconstructed in the remaining UM samples, we obtained a median of 64 [32.7-114.2] not statistically different from the median observed in CM (p=0.11, Mann-Whitney U-test). *Pf*EMP1 predicted domains for each patient, associated with expression value in FPKM are in supplemental table 2.

**Figure 4:**
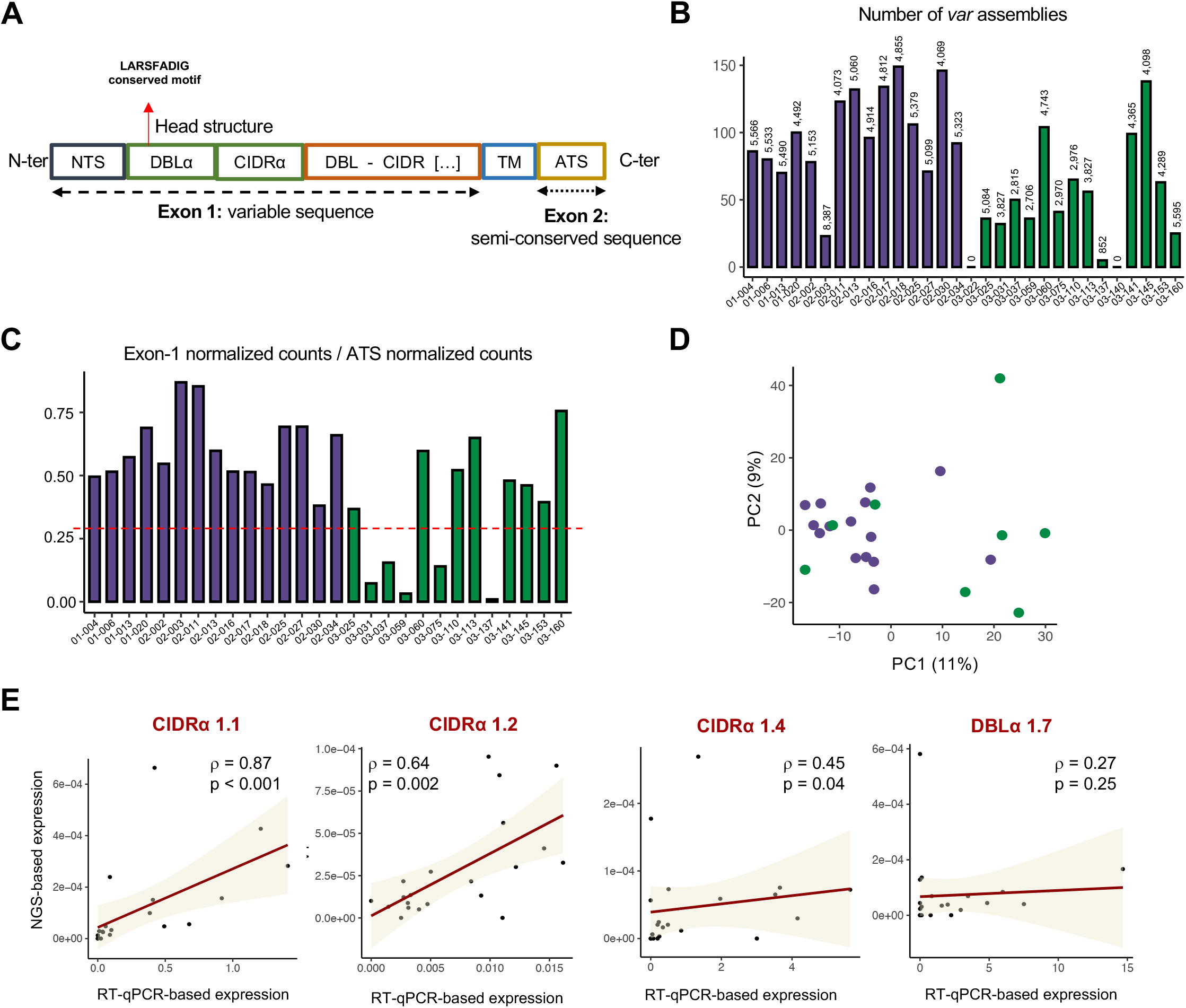
*Var* genes assembly and *var* genes domains expression. **A/** *Pf*EMP1 schematic organization. **B**/ *var* genes assembly statistics. The barplot represents the number of *var* assemblies for each patient with at least one predicted domain. The value at the top of each bar is N50 assembly statistic. **C**/ Proportion of assembled *var* genes in each sample. We compared the number of reads mapped to exon 1 and ATS to assess the quality of reconstructions. **D/** PCA based on *var* genes domains expression. **E**/ Correlation of NGS and RT-qPCR expression of *var* genes subdomains. Correlation was calculated with Spearman rank test.

In these isolates, sixteen full length *Pf*EMP1 sequences were obtained (8 CM and 2 UM). A search for the ICAM-1 binding motif showed the pattern in 12/15 CM and 5/8 UM isolates, confirmed by previous RT-qPCR assay (17). The pattern was concordantly not found by the two techniques in 2 CM and 3 UM and for 1 CM, the motif was found by RT-qPCR and not by RNA-seq. Regarding the PfEMP-1 protein structure, the DC8 domain (DBLα2-CIDRα1.1/8-DBLβ12) was found in 7/15 CM and 2/8 UM, and DC13 (DBLα1.7-CIDRα1.4) was found in 7/15 CM and 3/8 UM respectively. We considered that we have too few values in each clinical group to be able to compare the expression values. However, we performed PCA based on domains expression and noticed a partial grouping of CM isolates (Figure 4D). The correlation between NGS-based and RT-qPCR-based *Pf*EMP1 subdomain expression (17) showed a moderate (CIDRα1.4 ρ=0.45, p=0.04, Spearman rank test) to strong (CIDRα1.1 ρ=0.87 p<0.001 and CIDRα1.2 ρ=0.64 p=0.002; Spearman rank test) positive correlation (figure 4E).

### *var* genes abundance is higher in CM

We measured *var* genes expression using RNA-seq data, based on the ATS domain. Expression was normalized accounting for parasitaemia and IDC stage proportions. We found a 1.8-fold-increase in CM group (p=0.02, Mann-Whitney U-test). We validated these results through three independent RT-qPCR experiments, targeting both DBLα and ATS domain (see methods section, Figure 5, supplemental data 8). We found a 3.2-fold increase in CM with ATS-1 primers (p<0.001, Mann-Whitney U-test) and a 2.4-fold increase with ATS-2 primers (19) (p<0.001, Mann-Whitney U-test). A moderate to strong correlation (ρ>0.4, p≤0.05, Spearman rank test) was found between the RT-qPCR and NGS expression (figure 5A). In the additional group, we found a tendency of deregulation with ATS-1 primers (p=0.08, Mann-Whitney U-test) and we confirmed the difference with ATS-2 (p=0.03, Mann-Whitney U-test). As for DBLα primers (12), we observed a 2.2-fold increase in the sequenced group (p=0.09, Mann-Whitney U-test) but did not confirm the difference in the additional group (p=0.7, Mann-Whitney U-test) (Figure 5B). Then, we focused on the expression associated with assemblies’ sequences. We compared the expression data (RPKM normalized for developmental stage) of the most expressed assembly in each patient (minimum NTS-DBLα-CIDRα + 1 domain) and found a higher expression in CM, by a factor of 2.2 (p=0.007, Mann-Whitney U-test).

**Figure 5:**
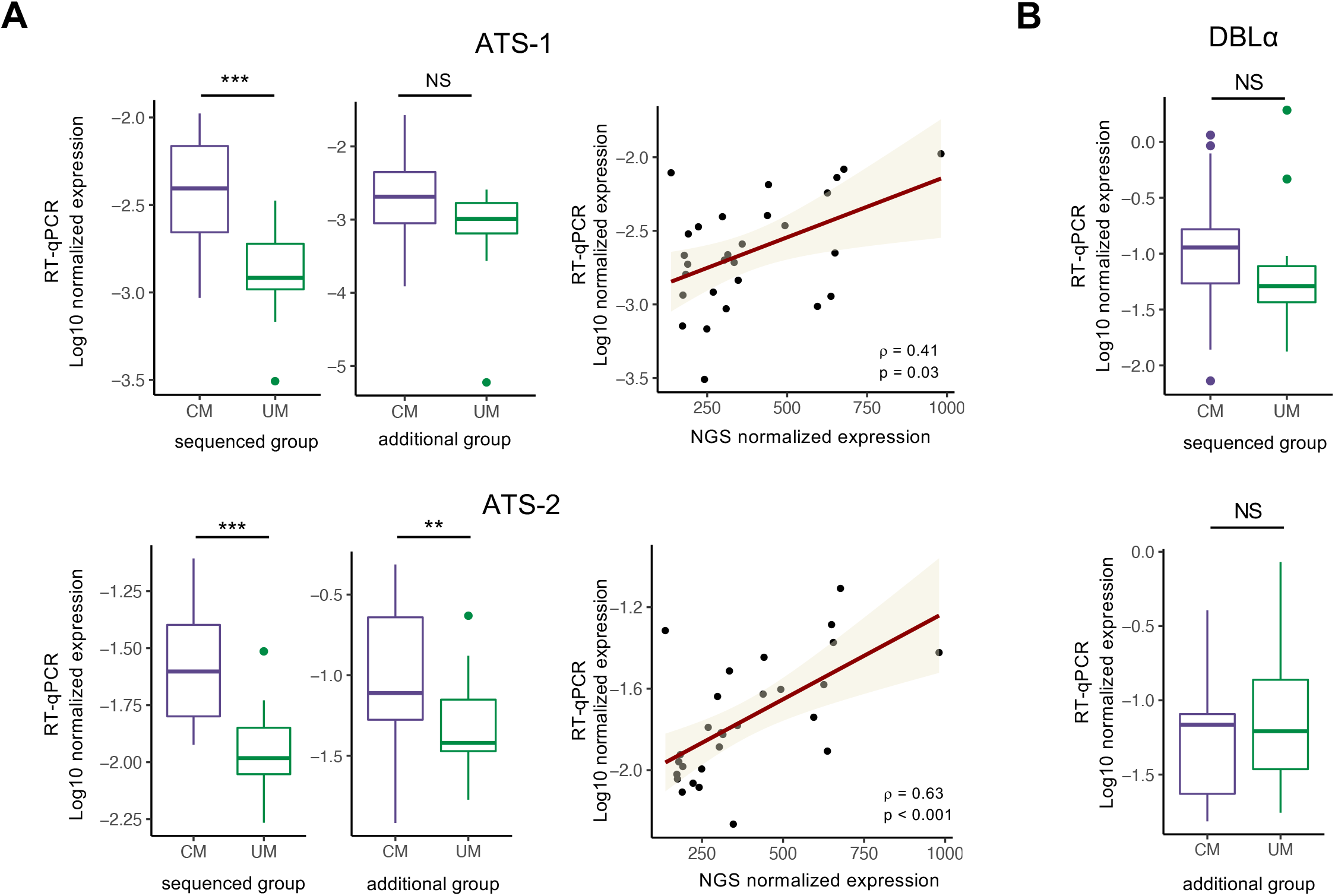
*Var* genes abundance measurement. **A/** *Var* gene expression based on ATS domain. Expression was measured by RT-qPCR using two primers sets. NGS and RT-qPCR ATS-based expression were correlated with Spearman rank test (Spearman rank correlation coefficient ρ and p-value are shown). **B/** *Var* gene expression based on DBLα domain, measured by RT-qPCR.

## Discussion

CM pathophysiology is related to *P. falciparum* ability to adhere to specific receptors in host brain capillaries (2,3), but host and parasite factors leading to neuroinflammation in CM are not yet fully elucidated. Although many recent studies have focused on host transcriptome signature associated with CM (20–22), the gene expression profile of *P. falciparum* isolated from CM remains largely unknown. In this context, we analyzed the gene expression of 15 CM and 15 UM *Plasmodium falciparum* isolates from a Beninese children population, to better understand the parasitic factors associated with CM.

Based on a published pipeline (5) which integrates data acquired from 3D7 synchronized strain (23), an increased proportion of circulating ring stages parasites was measured in CM isolates, suggesting a decreased circulation time compared to UM. Ring stage transcriptional signature was already associated with SM vs UM, as for acute malaria vs asymptomatic form (5–7). A recent retrospective study including datasets with various disease severity, including microarray data from Malawian children presenting with CM (24), also confirmed that lower circulating parasites hpi was correlated with disease severity (8).

Andrade *et al* predicted through a mathematical model that cytoadhesion alone may explain the parasites age difference observed in febrile compared to asymptomatic malaria isolates in a cohort of Malian individuals. Previous *in vitro* cytoadherence experiments using fresh parasites isolates from our cohort, showed that CM isolates exhibited a higher cytoadherence level on a human brain endothelial cell line (Hbec-5i) (17), which is a frequently used cellular model of CM (25,26). However, adherence levels differences should be confirmed through more efficient measures, as binding assay under flow conditions (18), or using a 3D human brain microvessel model (27).

Consistent with the hypothesis of an iE increased adherence capacity in CM, we found an overall upregulated expression of *var* genes in CM. For ATS domain-based measure, an upregulated expression of a factor 1.8 to 3.2 was established by both RNA-seq and RT-qPCR assay. For DBLα domain-based *var* genes expression, a non-significant 2.2-fold increase (p=0.1) in CM isolates was measured by RT-qPCR. Since DBLα sequences are more diverse (11), the nature of the corresponding primers (including degenerated bases) (12) may impact the quality of detection and subsequent quantification. By comparing the most expressed *var* gene in each sample, a trend of increased expression was reported in febrile malaria compared to asymptomatic infections (7). Tonkin-Hill *et al* did not measure a difference in *var* genes abundance in SM compared to UM, on the basis of *de novo* assemblies (5).

The *var* genes assemblies’ quality obtained in UM was not sufficient to perform domain expression comparison. A satisfactory reconstruction rate was measured in only half of the UM patients, significantly reducing the power of the DE analysis. These decreased reconstructions qualities were directly linked to the patients lower parasitaemia thus affecting the depth of the sequencing. RNA-seq data assemblies quality showed that the RT-qPCR approach we previously applied to our cohort (17) was more powerful. A way to have more reliable *var* genes domains expression data would have been to use the DBLα-tag approach (6,12). *var* genes transcripts may also be amplified through long-range PCR assays and analyzed by Illumina deep sequencing or PacBio long-read sequencing (28). The advantage of a long-read sequencing is to overcome *in silico* assembly of *var* genes. Although DE analysis was not performed, the PCA based on *var* genes domains expression showed a grouping of CM, which may reflect a more restricted *var* genes repertoire. Indeed, CM isolates express *var* genes domains allowing specific adhesion to EPCR, which we previously confirmed in our cohort, and ICAM-1 (12,13,17,18). Regarding the expression of the DBLβ1/3 ICAM-1 binding motif (14), the comparison with the RT-qPCR results showed very concordant results, finding an expression in both CM and UM, still questioning the implication of this motif in *Pf*EMP1 binding to ICAM-1 receptor (17).

In addition to iE increased adherence capacity mainly mediated by *Pf*EMP1, an efficient immune response against iE surface proteins, may also affect adhesion (9). We did not confirm this hypothesis by total antibodies titers, or by anti-*P. falciparum* IgG measured in the children plasma. However, we previously measured higher IgG levels against DBLβ3 and CIDRα1.4-DBLβ3 domains in the UM children from our cohort (17). Further experiments including more recombinant proteins from specific domains of *Pf*EMP1 could increase the accuracy of the measurement (29). In addition, binding inhibition assays could help to understand more precisely the influence of the immune system in iE cytoadhesion mechanisms in further studies (30-32). However, humoral immunity is not the only host protection against malaria infection and the role of complement component 1s, modulating iE adherence, has recently been demonstrated (33).

The decreased circulation time observed in CM may result in iE avoidance of splenic clearance, leading to higher parasitaemia (7). Interestingly, we pointed out a greater parasitaemia in CM in our sequenced group. A correlation between decreased circulation time and increased parasitaemia was retrospectively demonstrated from different studies, by Thomson-Luque and colleagues (8). However, this result should be modulated as no difference in parasitaemia was demonstrated in the additional group, whereas more ring stage circulating parasites was found in CM.

As adhesion capacity does not fully explain CM pathophysiology, we then focused on all-genes expression, excluding VSAs, to decipher CM specificities. After normalization accounting for parasite developmental stage, gene expression profile was a signature of malaria severity. PCA showed a total segregation of CM and UM clinical presentation, meaning the parasite “core” transcriptome is also involved in CM pathophysiology. More than 500 genes were deregulated in the CM group compared to the UM isolates. The results were confirmed by RT-qPCR for 8 genes (4 upregulated in CM and 4 downregulated) with a strong correlation of gene expression measured by RNA-seq and RT-qPCR. Importantly, gene expression was also assessed in the additional group by RT-qPCR and deregulation was confirmed for the four upregulated genes in CM but not the downregulated genes. In fact, UM is a heterogeneous group, which may bring some variability in the differential measurement of gene expression.

Focusing on differentially expressed genes, DNA and protein metabolism pathways were dysregulated in CM samples, which reflects the overall gene expression dysregulation. Transcripts in glucose- and pyruvate-associated pathways were dysregulated, but we could not correlate our results with a metabolically quiescent phenotype in CM, as previously reported in SM (5). Plasmepsin IX (PMIX, PF3D7_1430200) and Plasmepsin X (PMX, PF3D7_0808200) proteins, involved in the GO pathway entry into host, were upregulated in CM. PMIX is essential for the parasite invasion of erythrocyte and PMX is essential for both exit and invasion. These two proteins are druggable mediators, implicated in both entry and exit from host (34). Erythrocyte binding antigen-175 (eba-175, PF3D7_0731500), a receptor having a key role in the binding between merozoite and erythrocyte (35), was also upregulated. Apical membrane antigen 1 (AMA-1, PF3D7_1133400) gene expression was upregulated in CM and was in the top 20 deregulated genes. The corresponding protein is implicated in erythrocyte invasion (36). AMA-1 has a crucial role in malaria pathophysiology, and a malaria vaccine targeting this protein was already produced, and tested (37). Eight other proteins implicated in entry into host were found upregulated. It could reflect an increased invasion capacity of CM isolates parasites, that may also promote the greater parasitaemia measured in the CM.

Regarding pathways implicated in cytoadhesion and in *Pf*EMP1 regulation, we also found numerous upregulated genes in CM. PHISTa was the most upregulated gene in CM and PHISTb gene (PF3D7_0424600) was also upregulated. Several members of the PHIST family have been described to act as an anchor for *Pf*EMP1 by binding to the ATS domain (38). This protein has already been characterized as overexpressed in pregnancy associated malaria (39,40). CLAG 9 gene was up-regulated in CM and the corresponding protein was identified as essential for VAR2CSA presentation, which means that it plays a major role in addressing *Pf*EMP1 to the membrane (41). Other proteins implicated in *Pf*EMP1 regulation were dysregulated as SET9, 10 and EXP2, 3 involved in the regulation of *var* genes expression and the export of *Pf*EMP1 through the parasitophorous vacuole (PV) membrane (42).

In summary, 15 CM and 15 UM *P. falciparum* transcriptomes were successfully sequenced. An increased ring stage transcriptomic signature was identified in CM circulating parasites compared to UM, that may result from an increased parasites adherence capacity. Indeed, a higher adherence was measured on CM iE, for two different cell lines. Consistent with this result, *var* genes were overexpressed in CM. *var* genes domains expression was more restricted in CM isolates compared to UM, reflecting the specific binding to receptors in host brain endothelium capillaries. However, ICAM-1 binding motif presence was confirmed in both CM and UM, still questioning its role in *Pf*EMP1 adhesion to ICAM-1 receptor. We also found numerous upregulated genes involved in entry into host pathway, which could reflect an increased invasion capacity of CM isolates. Deregulated genes involved in adhesion support the hypothesis of a stronger CM adhesion. UM parasites increased circulation time may also be due to a more efficient immune response in UM, which we could not demonstrate on our cohort. To conclude, parasitic and host factors leading to neuroinflammation in CM should be further investigate and particularly factors influencing parasite decreased circulation time. Difference in term of intrinsic parasite adhesion capacity linked to total *var* genes overexpression and increased invasion of erythrocytes should be deeply explored.

## Material et methods

### Samples recruitment

Ethical clearance was obtained from Comité National d’Ethique pour la recherche en santé au Bénin (N°67/MS/DC/SGM/DRFMT/CNERS/SA; 17 October 2017) and by Comité consultative de déontologie et d’éthique of Institut de Recherche pour le Développement (IRD; 24 October 2017). Children under 6 years old presenting with CM or UM were recruited from December 2017 to November 2018 in South Benin. Malaria infection was diagnosed with positive *P. falciparum* thin or thick blood smear. For CM, a Blantyre score <= 2 was required. A patient was included as UM if he displayed a parasitaemia between 1000 to 5.10^5^ parasites/μl and no clinical or biological signs of severe malaria. Only patients with *P. falciparum* mono-infection were considered. The recruitment of samples has been detailed previously (16) (17).

### Measurement of anti-*P. falciparum* antibodies

Detection and quantification of total antibodies against *P. falciparum* was performed to evaluate previous exposure to *Plasmodium sp*. IgG/A/M anti-plasmodial antibodies were detected and quantified by indirect immunofluorescence assay using whole schizonts of the 3D7 *P. falciparum* strain as crude antigens, and fluoresceine linked anti-human IgG/A/M (Biorad^®^; Hercules, California, USA) as conjugate. Quantification of plasmatic antibody concentration was estimated by serologic titers after a four-fold dilution series of plasma from 1:16 to 1:4096 (43). Slides were examined under blinded conditions by two experienced microscopists. In case of discordant result, a third microscopist observed in a blinded manner the slide to arbitrate. The validation of the method determined the threshold of positivity at a titer of (1:64). Antibody titers from (1:64) to (1:1024) are usually observed at malaria remission. Titers over or equal to (1:4096) are unusual. For statistical analysis, antibody titers were classified into three groups: negative (or 0) corresponding to titers < 1:64; positive, corresponding to titers 1:64, 1:256 and 1:1024 and strongly positive, corresponding to titers ≥ 1:4096 (43). To measure the difference between the clinical phenotypes, we considered the titers as a qualitative variable and used χ^2^ test.

IgG were also detected and quantified by flow cytometry on 3D7 *P. falciparum* strain. Briefly, mature parasitic forms cultivated at 37°C in a controlled atmosphere at 95% humidity with a gas mixture of 5% CO_2_, 10% O_2_ and 85% N_2_, in healthy red blood cells of group O ^+^ (EFS, Bichat Hospital, France) and RPMI Gibco^®^ 1640 medium (Life technologies^®^; Carlsbad, California, USA) supplemented with 10% human serum from group AB (Biowest®, Nuaillé, France), HEPES 0.5% (Sigma-Aldrich®, Saint Louis, Missouri, USA) and NaHCO_3_ (Sigma-Aldrich®) at 5% hematocrit were enriched by MACS and the concentrated parasite suspension was then adjusted at 10,000,000 iE/ml and 20% of parasitaemia. Patient plasma was brought into contact with the iE suspension and secondary revealed, after wash, by goat IgG anti human IgG Fc labelled with phycoerythrin. SYTO™ 61 label (Thermo Fisher Scientific) was also used for parasite label in iE. Measurements were performed after a count of 10,000 cells in duplicate. As negative controls, non-iE and plasma from unexposed malaria patient were used.

### Samples’ preparation and RNA sequencing

*P. falciparum* RNA was extracted from whole blood preserved at −80°C in TRIzol LS reagent (Life Technologies) with phenol-chloroform protocol and purified using RNeasy minikit (Qiagen). NanoDrop 2000c (Thermo Fisher Scientific) and Agilent RNA 6000 Pico Kit (Agilent Biotechnologies) were used to assess RNA quality, concentration, and contamination (detailed in supplemental data 1). Samples with both RNA Integrity Number (RIN) >= 6 and at least 100 ng of total RNA available were retained for sequencing. Based on these criteria, we kept 16 CM and 21 UM. TruSeq Stranded mRNA protocol (Illumina, California, U.S.A) was used for library preparation and sequencing was performed after qPCR quantification on a NovaSeq 6000 (Illumina) system, using 2*100 cycles (paired-end reads, 100 nucleotides) and aiming to obtain about 20M clusters per sample (Institut Curie, ICGex - NGS platform).

### Gene expression analysis

We used the scripts published by Tonkin-Hill *et al* (https://github.com/gtonkinhill/falciparum_transcriptome_manuscript) (5). Sequencing quality was assessed with FastQC v0.11.9 (44). Raw reads were mapped on both *Plasmodium falciparum* (3D7 strain, PlasmoDB v49) and human (hg38 UCSC genome browser) reference genomes with Hisat2 v2.2 (45). Aligned reads were assigned to *P. falciparum* annotated features from PlasmoDB-49_Pfalciparum3D7.gff with Feature-counts (46). VSA were kept out, as we cannot rely on reference genome. We excluded samples below a minimal cutoff of four million reads mapped to the parasite genome. Parasites’ blood stages proportions in each sample were evaluated as described in Tonkin-Hill *et al* (5), in which data acquired from synchronized strains of 3D7 are used (23). In the following analysis, ring stage corresponds to 8 hours post infection (hpi), early trophozoite at 19 hpi, late trophozoite at 30 hpi and schizont at 42 hpi. Counts were normalized accounting for both the library size and the parasite’s stage proportions, then differential expression analysis was performed using the R Limma Voom method (47) (5). EdgeR package (48) was used to import and normalize data. We consider a gene as differentially expressed for p-adj <= 0.05 (Benjamini-Hochberg) and a Fold-Change (FC) => |1.5|.

### Parasite developmental stage determination by microscopy

The parasites’ stage proportions were evaluated by optical microscopy on thin blood smears, using previous classification by Silamut *et al* (3). To best correspond to the classification determined from the high-throughput sequencing data, the stages were evaluated as follows: the ring stage corresponds to parasites between 0 and 16 hpi, the early trophozoites correspond to parasites between 16 and 30 hpi, late trophozoites 30 to 38 hpi and schizonts 38 to 48 hpi.

### Additional group

We designed an additional group to reconfirm results from the sequenced group. Samples were chosen randomly and based on the possibility of establishing the proportions of stages by reading the thin blood smears. We kept 15 CM and 15 UM to have the same number of patients as in the sequenced group.

### RT-qPCR validations of the all-gene expression bioinformatic analysis

Primers were designed using Primer3Plus online tool (49). RT-qPCR were performed using the SensiFAST Sybr No-ROX kit (Bioline) on the Rotor-Gene qPCR system (Qiagen) following these PCR conditions: 95°C for 2 s; 40 cycles of 95°C for 5 s, 60 to 62°C for 10 s, and 72°C for 10 s; and a dissociation phase from 60°C to 90°C. Gene levels transcripts were evaluated with serial dilutions of plasmids (Genscript). Results were normalized with a house-keeping gene (seryl-tRNA synthetase gene expression (50)) and the parasite’s stage proportions (RNA-seq data (23) were used to calculate a normalization factor based on calculated proportions for the sequenced group or readed proportions for the additional group). We used human and 3D7 cDNA as negative and positive controls at each RT-qPCR run.

### Differentially expressed (DE) genes analysis: GSEA and Protein-protein interaction (PPI) network

The Gene Ontology (GO) (51) and KEGG pathway (52) enrichment analyzes were performed by applying a hypergeometric test on the PlasmoDB GO annotation file (PlasmoDB-49_Pfalciparum3D7_GO.gaf) and on the KEGG annotations downloaded from PlasmoDB. Enriched pathways with p-value less than 0.05 were considered for further analysis. The PPI network was generated from DE genes in the STRING online site (v. 11.5) (53). Settings were set to high confidence (0.7) interaction score and co-expression interactions source. Then, cluster analysis was performed in Cytoscape 3.8.2 (54) with MCode application v2.0 (55).

We kept the co-expression interactions with a high degree of confidence. Co-expression is based on gene-by-gene correlation tests across many gene expression datasets (using both transcriptome and proteome measurements).

### *var* genes transcripts assembly

Var transcripts were assembled *de novo* from RNA-seq reads using a published pipeline (https://github.com/PapenfussLab/assemble_var) (5). Briefly, reads were mapped against *P. falciparum* 3D7 (PlasmoDB) and human (hg38, UCSC genome browser) reference genomes. Reads unmapped and mapped against 3D7 *var* genes were extracted and used for assembly with oases and cap3. After, contigs < 500 nucleotides length were discarded. *P. falciparum* reads from each isolate were aligned with R subreads against their own reconstructed var transcripts. Then, the redundant contigs were filtered out (overlap = 0.7 and identity = 95%) and finally the remaining contigs were translated into proteins. The largest open reading frame was kept. Protein sequences less than 130 amino acids were excluded. The positioning of each domain (NTS - DBLα/β/δ/γ/ε/ζ - CIDRα/β/δ/γ -ATS) was performed using Vardom online tool (11). Sequences predicted as not containing at least one full-length domain were not considered for downstream analysis. Then, the nature of the subdomain (ie DBLα1.1-2-3/2) was assessed by performing an alignment with MAFFT (56) against the Vardom database (11). We then manually filtered out the aberrant domains enchainments. Assemblies’ quality was assessed by comparing the number of reads mapping to exon 1 assemblies and ATS domain. We can estimate that if we have reconstructed the total set of var transcripts, we will have an equivalent number of aligned reads. As ATS is a semi-conserved domain, to map the maximum number of reads, we relied on both assembled ATS and ATS published from published databases (Vardom seven reference genomes (12) and varDB (57)). The number of reads were normalized by exon 1 or ATS domain average size in 3D7 reference.

### *var* genes expression measured by RNA-seq and RT-qPCR

Raw reads were aligned with Hisat2 v2.2.1 (45) to a database composed of ATS domains from Vardom’s 7 reference genomes (11), varDB (57) and RNA-seq assembly. Counts were normalized accounting for both the library size and the parasite’s stage proportions (RNA-seq data (23) were used to calculate a normalization factor based on calculated proportions). *var* genes expression was confirmed through three independent RT-qPCR. We first used the DBLα primers developed by Lavstsen *et al* (12) for DBLα tag. We also developed primers targeting the *var*-ATS using varDB (57) ATS sequences, primers sequences and ATS-targeting statistics are given in supplemental data 9 (ATS-2). The two preceding qPCR were performed as previously described with the SensiFAST Sybr No-ROX kit (Bioline) (see the paragraph above for more precision). Finally, we also performed the qPCR-Taqman developed by Hofmann *et al* targeting the ATS part of the *var* (19) (ATS-1) on a Viia7 (Applied Biosystems) with a slightly modified protocol. Briefly, 1 μl of diluted cDNA was amplified with 250 nM of MGB-probe (Eurogentec), 400 nM of primer-fw and primer-rv, 10 μl of Luna Universal probe qPCR master mix (New England Biolabs, Ipswich, USA) in a final volume of 20 μl. The thermal profil used was as follows: 60s at 95°C, 45 cycles of 15 s at 95°C and 45 s at 55°C. Results were normalized as described above.

### Statistical analysis

Graphics and statistical analysis were realized using R software. In case of non-normality distribution, median values were calculated and presented with 10th and 90th percentile. Differences were evaluated with the Wilcoxon Mann Whitney nonparametric test. Proportions were compared with χ^2^ test. Correlations were evaluated with the Spearman correlation test, using ρ and p-value. P-values ≤ 0.05 were considered as significant.

## Data availability

RNA-seq data have been deposed in GEO repository GSE186820.

## Acknowledgments

We are grateful to all patients and their parents for participating in this study. We are thankful to the nurses of the CHU-MEL hospital (CHU-Mère et Enfant de la Lagune) and Hôpital de zone de Calavi for their help in the collection of samples. We also thank the health center of So-Ava. We acknowledge Antoine Claessens and all his team, Prince Nyarko, Marc-Antoine Guéry and Camille Cohen for their help and counselling regarding data analysis. High-throughput sequencing was performed by the ICGex NGS platform of the Institut Curie supported by the grants ANR-10-EQPX-03 (Equipex) and ANR-10-INBS-09-08 (France Génomique Consortium) from the Agence Nationale de la Recherche (“Investissements d’Avenir” program), by the ITMO-Cancer Aviesan (Plan Cancer III) and by the SiRIC-Curie program (SiRIC Grant INCa-DGOS-465 and INCa-DGOS-Inserm_12554).

## Contributors

E.G.: designed the experiments; performed the experiments; bioinformatic methodology; data analysis; wrote the article. J.F.: designed the experiments; performed the experiments; revised the manuscript. V.J.: designed the experiments; performed the experiments; revised the manuscript. C.K.: designed experiments; revised the manuscript. B.V.: performed the experiments. L.H.: performed the experiments. J.F.F.: supervised patients’ recruitment; clinical analysis. A.A.: supervised patients’ recruitment. S.H.: supervised clinical analysis. M.C.: revised the manuscript. N.A.: supervised the experiments; performed the experiments; revised the manuscript. O.T.: supervised data analysis; revised the manuscript. G.I.B.: designed the study; supervised the experiments; supervised data analysis; wrote the article.

## NeuroCM group

Jules Alao (Pediatric Department, Calavi Hospital, Calavi, Benin); Dissou Affolabi (Pediatric Department, Calavi Hospital, Calavi, Benin); Bibiane Biokou (Pediatric Department, Mother and Child University and Hospital Center (CHUMEL), Cotonou, Benin); Jean-Eudes Degbelo (Institut de Recherche Clinique du Bénin (IRCB), Calavi, Benin); Philippe Deloron (MERIT UMR 261, IRD, Université de Paris, Paris, France); Latifou Dramane (Institut de Recherche Clinique du Bénin (IRCB), Abomey Calavi, Benin); Sayeh Jafari-Guemouri (MERIT UMR 261, IRD, Université de Paris, Paris, France); Anaïs Labrunie (NET, INSERM, Université de Limoges, Limoges, France); Yélé Ladipo (Pediatric Department, Mother and Child University and Hospital Center (CHUMEL), Cotonou, Benin); Thomas Lathiere (Ophtalmology department, Limoges University Hospital, Limoges, France); Achille Massougbodji (Institut de Recherche Clinique du Bénin (IRCB), Abomey Calavi, Benin); Audrey Mowendabeka (Paediatric Department, Hopital de la Mère et de l’Enfant, Limoges, France); Jade Papin (MERIT UMR 261, IRD, Université de Paris, Paris, France); Bernard Pipy (PHARMADEV, Université de Toulouse, IRD, UPS, France); Pierre-Marie Preux (NET, INSERM, Université de Limoges, Limoges, France); Marie Raymondeau (NET, INSERM, Université de Limoges, Limoges, France); Jade Royo (PHARMADEV, Université de Toulouse, IRD, UPS, France); Darius Sossou (Institut de Recherche Clinique du Bénin (IRCB), Abomey Calavi, Benin); Brigitte Techer (MERIT UMR 261, IRD, Université de Paris, Paris, France).

## Supplement

**Supplemental data 1: RNA quality.** Nanodrop 260/280, 260/230 ratios and rRNA ratio and RIN values from Bioanalyzer Pico Chip. Bioanalyzer Pico Chip Electropherograms.

**Supplemental data 2: Mapping statistics.** Total sequenced reads and corresponding mapping to *P. falciparum* and human genomes are displayed.

**Supplemental data 3: Antibodies anti-*P. falciparum* measurement.** Quantification of total antibodies against *P. falciparum* iE with indirect fluorescence. IgG anti-VSA detection and quantification by flow cytometry.

**Supplemental data 4: Parasites developmental blood stages.** Parasites stages proportions for CM and UM from the sequenced group, measured from RNA-seq expression and microscopic reading of thin blood smears. Parasites stages proportions for CM and UM from additional group, evaluated by microscopy.

**Supplemental data 5: RT-qPCR validations.** For each gene, we show median with 10^th^ and 90^th^ percentile in CM and UM clinical group. The p-values were calculated with the Mann-Whitney U-test. For the sequenced group, the correlation between the normalized expression values measured by RT-qPCR and RNA-seq was represented through the regression line and tested by the Spearman rank test (Spearman rank correlation coefficient ρ and p-value are given).

**Supplemental data 6: GSEA - GO biological process and KEGG pathways.** Deregulated genes belonging to each GO category or KEGG pathway. Genes upregulated in CM compared to UM are colored in red, genes downregulated in CM in blue.

**Supplemental data 7: MCode clusters from PPI network.** Deregulated genes belonging to each MCode cluster. Genes upregulated in CM compared to UM are colored in red, genes downregulated in CM in blue.

**Supplemental data 8: *var* genes expression measurement by RT-qPCR.** For each primers set, we show median with 10^th^ and 90^th^ percentile in CM and UM clinical group. The p-values were calculated with the Mann-Whitney U-test.

**Supplemental data 9: *var* ATS-2 primers.** Newly designed primer sequences and mapping statistics on Vardom and VarDB databases.

**Supplemental table 1: Differentially expressed genes excluding VSAs between CM and UM.**

**Supplemental table 2: *var* genes transcripts assemblies for each patient**

