## Supplemental data for "Transcriptome analysis of *Plasmodium falciparum* isolates from Benin reveals specific gene expression associated with cerebral malaria"

#### Supplemental data 1: RNA quality

| Sample | Nanodrop | | Bioanalyzer profil Pico Chip | |
|---|---|---|---|---|
| | **Ratio 260/280** | **Ratio 260/230** | **rRNA Ratio [28s/18s]** | **RIN** |
| 01-004 | 1.98 | 1.26 | 0.2 | 7.6 |
| 01-006 | 2.25 | 2.3 | 1.8 | 8.7 |
| 01-013 | 2.09 | 0.93 | N/A | 8.5 |
| 01-020 | 2.26 | 0.24 | N/A | 7.5 |
| 01-033 | 2.05 | 2.13 | 1.3 | 7.1 |
| 02-002 | 2.24 | 2.24 | N/A | 6.9 |
| 02-003 | 1.89 | 0.9 | 1.2 | 7.2 |
| 02-011 | 2.2 | 0.42 | 0.9 | 8 |
| 02-013 | 2.19 | 1.96 | 1.3 | 7.2 |
| 02-016 | 2.2 | 1.65 | 0.6 | 6.6 |
| 02-017 | 2.22 | 2.32 | 1.7 | 7.3 |
| 02-018 | 1.97 | 1.35 | 0.9 | 9.6 |
| 02-025 | 2.2 | 1.66 | 1.1 | 8.5 |
| 02-027 | 2.21 | 1.98 | 0.2 | 7 |
| 02-030 | 2.19 | 1.66 | N/A | 8 |
| 02-034 | 2.22 | 1.79 | 0.8 | 9 |
| 03-022 | 1.73 | 0.84 | 0.2 | 6.6 |
| 03-025 | 2.22 | 2.09 | 1.1 | 9 |
| 03-028 | 2.05 | 1.58 | 1.1 | 6.9 |
| 03-030 | 2.04 | 0.97 | 1.0 | 7.1 |
| 03-031 | 2.11 | 1.9 | 1.7 | 7.4 |
| 03-037 | 2.16 | 2.34 | N/A | 7.1 |
| 03-051 | 1.82 | 1.13 | 0.7 | 7.1 |
| 03-059 | 2.12 | 1.75 | 1.5 | 9.7 |
| 03-060 | 2.09 | 1.68 | N/A | 7.3 |
| 03-075 | 2.09 | 0.69 | N/A | 7.1 |
| 03-108 | 2.02 | 1.54 | 1.0 | 8.1 |
| 03-110 | 2.03 | 1.71 | N/A | 6.7 |
| 03-113 | 1.9 | 1.09 | 0.3 | 7.4 |
| 03-127 | 1.97 | 1.09 | 0.6 | 7 |
| 03-134 | 2.11 | 0.13 | 0.9 | 6.8 |
| 03-137 | 1.91 | 0.15 | N/A | 6.9 |
| 03-140 | 2.26 | 2.47 | 1.6 | 7.7 |
| 03-141 | 2.21 | 1.55 | 1.3 | 9.7 |
| 03-145 | 1.8 | 0.75 | 1.5 | 7.6 |
| 03-153 | 2.1 | 1.71 | 0.9 | 9.8 |
| 03-160 | 2.09 | 1.27 | 3.0 | 7.8 |

01-004

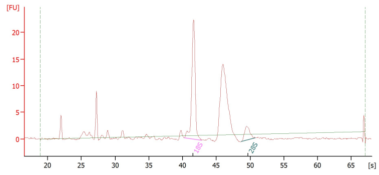

01-006

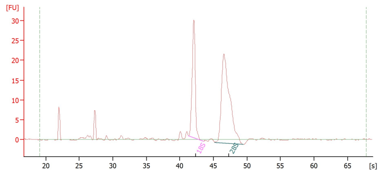

01-013

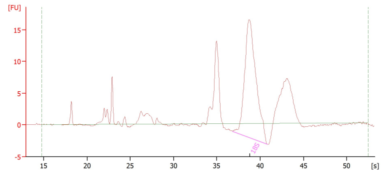

01-020

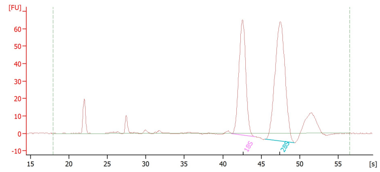

01-033

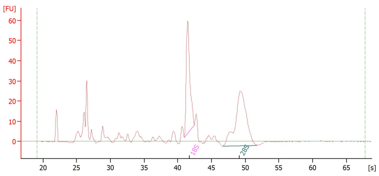

02-002

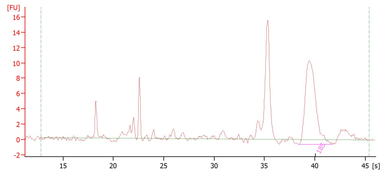

02-003

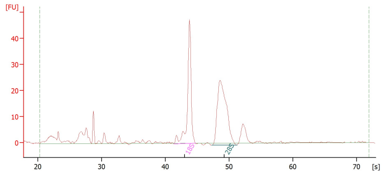

02-011

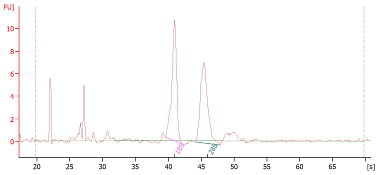

02-013

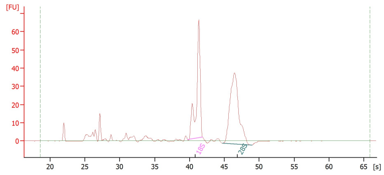

02-016

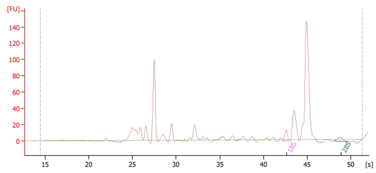

02-017

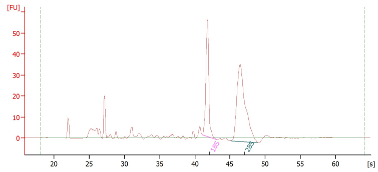

02-018

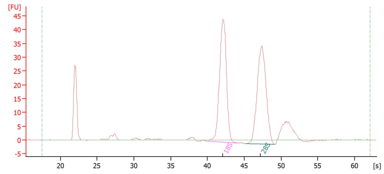

02-025

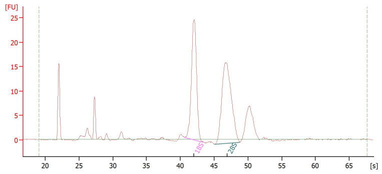

02-027

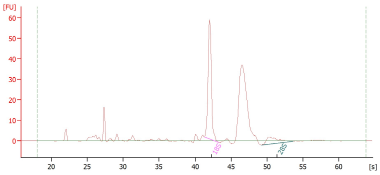

02-030

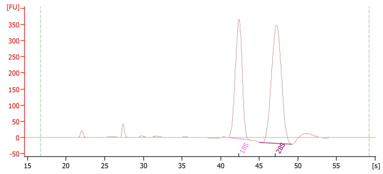

02-034

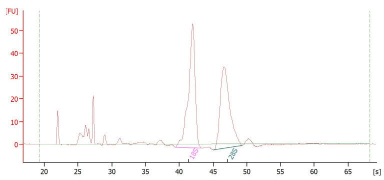

03-022

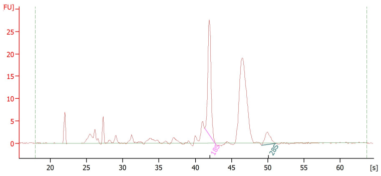

03-025

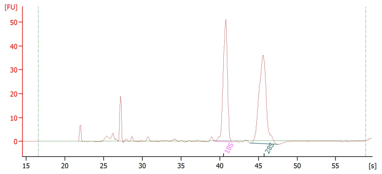

03-028

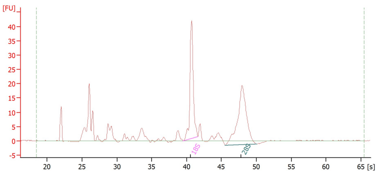

03-030

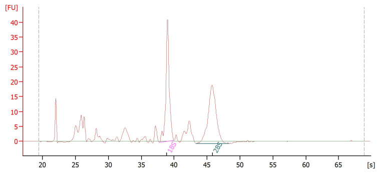

03-031

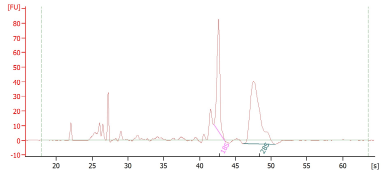

03-037

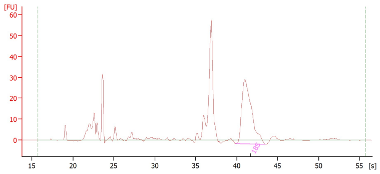

03-051

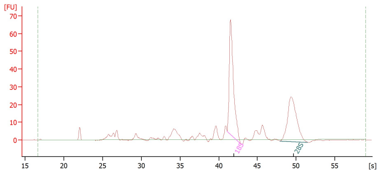

03-059

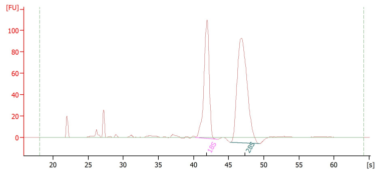

03-060

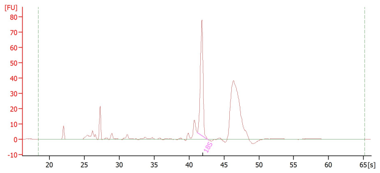

03-075

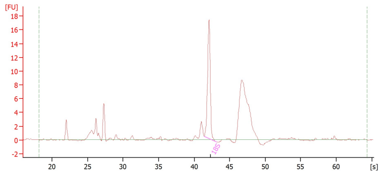

03-108

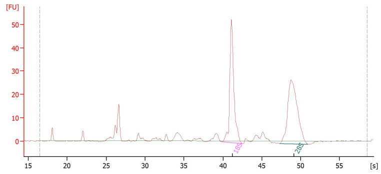

03-110

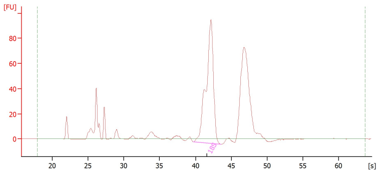

03-113

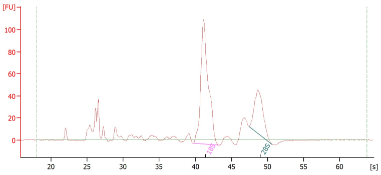

03-127

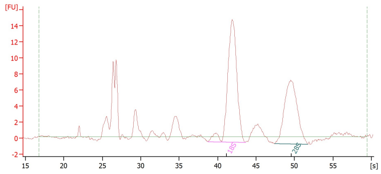

03-134

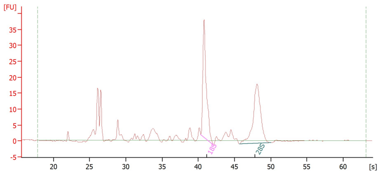

03-137

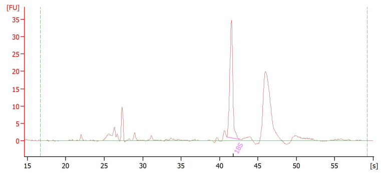

03-140

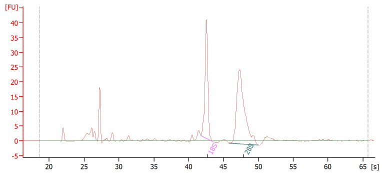

03-141

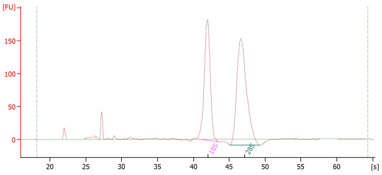

03-145

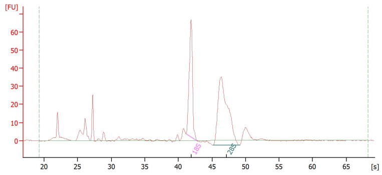

03-153

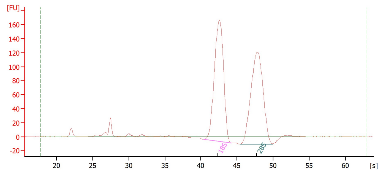

03-160

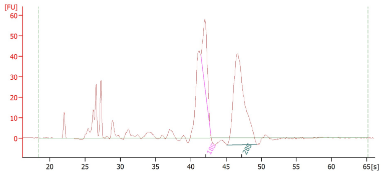

#### Supplemental data 2: Mapping statistics

| Sample | Total PE reads | Reads aligned on *P. falciparum** | Reads aligned on human* | % aligned on *P. falciparum* | Included |
|---|---|---|---|---|---|
| 01-004 | 29,460,649 | 25,013,688 | 1,806,283 | 84.91 | yes |
| 01-006 | 30,339,991 | 22,362,568 | 4,196,159 | 73.71 | yes |
| 01-013 | 19,726,822 | 15,530,598 | 2,581,835 | 78.72 | yes |
| 01-020 | 24,824,089 | 22,115,932 | 367,422 | 89.09 | yes |
| 01-033 | 24,725,082 | 842,203 | 21,956,993 | 3.41 | no |
| 02-002 | 29,967,330 | 23,760,342 | 3,414,561 | 79.29 | yes |
| 02-003 | 25,691,710 | 21,589,730 | 1,440,628 | 84.03 | yes |
| 02-011 | 20,387,129 | 13,886,245 | 3,629,496 | 68.11 | yes |
| 02-013 | 25,766,867 | 17,300,223 | 6,317,238 | 67.14 | yes |
| 02-016 | 34,260,826 | 28,218,798 | 2,673,820 | 82.36 | yes |
| 02-017 | 22,279,539 | 18,674,319 | 1,598,497 | 83.82 | yes |
| 02-018 | 24,899,632 | 13,481,147 | 9,448,237 | 54.14 | yes |
| 02-025 | 34,465,539 | 30,214,204 | 552,149 | 87.66 | yes |
| 02-027 | 21,078,309 | 17,703,933 | 1,482,800 | 83.99 | yes |
| 02-030 | 18,863,558 | 14,905,305 | 1,827,742 | 79.01 | yes |
| 02-034 | 40,413,108 | 30,939,185 | 5,556,226 | 76.56 | yes |
| 03-022 | 22,443,761 | 11,821,917 | 8,357,942 | 52.67 | yes |
| 03-025 | 22,891,522 | 19,206,253 | 1,534,413 | 83.90 | yes |
| 03-028 | 26,474,232 | 2,327,672 | 21,951,661 | 8.79 | no |
| 03-030 | 40,117,029 | 931,038 | 36,233,376 | 2.32 | no |
| 03-031 | 22,677,427 | 13,686,139 | 7,078,106 | 60.35 | yes |
| 03-037 | 25,984,157 | 18,574,199 | 5,313,743 | 71.48 | yes |
| 03-051 | 27,872,854 | 367,849 | 25,725,922 | 1.32 | no |
| 03-059 | 23,166,613 | 19,020,322 | 2,098,601 | 82.10 | yes |
| 03-060 | 22,842,008 | 18,788,367 | 2,057,835 | 82.25 | yes |
| 03-075 | 17,905,072 | 13,229,748 | 2,900,006 | 73.89 | yes |
| 03-108 | 38,637,117 | 13,134 | 34,668,977 | 0.03 | no |
| 03-110 | 25,517,266 | 15,103,898 | 8,062,731 | 59.19 | yes |
| 03-113 | 23,284,009 | 4,198,073 | 16,458,818 | 18.03 | yes |
| 03-127 | 41,860,202 | 611,208 | 36,757,232 | 1.46 | no |
| 03-134 | 28,279,149 | 106,671 | 26,422,856 | 0.38 | no |
| 03-137 | 36,010,543 | 27,498,890 | 4,726,525 | 76.36 | yes |
| 03-140 | 19,892,897 | 16,525,296 | 1,709,248 | 83.07 | yes |
| 03-141 | 18,449,529 | 15,826,946 | 827,848 | 85.78 | yes |
| 03-145 | 19,143,568 | 15,935,664 | 1,057,201 | 83.24 | yes |
| 03-153 | 18,790,453 | 11,106,406 | 5,732,515 | 59.10 | yes |
| 03-160 | 19,916,399 | 9,969,847 | 7,765,547 | 50.06 | yes |

*PE reads aligned concordantly

#### Supplemental data 3: Antibodies anti-*P. falciparum* measurement

- **Quantification of total antibodies against *P. falciparum* iE with indirect fluorescence**

| sample | clinical group | Antibody titer | immunological statut |
|---|---|---|---|
| 01-004 | CM | 4096 | strongly positive |
| 01-006 | CM | 4096 | strongly positive |
| 01-013 | CM | 1024 | strongly positive |
| 01-020 | CM | 256 | positive |
| 02-002 | CM | 4096 | strongly positive |
| 02-003 | CM | 256 | positive |
| 02-011 | CM | 4096 | strongly positive |
| 02-013 | CM | 1024 | strongly positive |
| 02-016 | CM | 1024 | strongly positive |
| 02-017 | CM | 256 | positive |
| 02-018 | CM | 1024 | strongly positive |
| 02-025 | CM | 1024 | strongly positive |
| 02-027 | CM | 1024 | strongly positive |
| 02-030 | CM | 256 | positive |
| 02-034 | CM | 4096 | strongly positive |
| 03-022 | UM | 1024 | strongly positive |
| 03-025 | UM | 4096 | strongly positive |
| 03-031 | UM | 4096 | strongly positive |
| 03-037 | UM | 4096 | strongly positive |
| 03-039 | UM | 256 | positive |
| 03-060 | UM | 1024 | strongly positive |
| 03-075 | UM | 1024 | strongly positive |
| 03-110 | UM | 1024 | strongly positive |
| 03-113 | UM | 256 | positive |
| 03-137 | UM | 64 | positive |
| 03-140 | UM | 256 | positive |
| 03-141 | UM | 64 | positive |
| 03-145 | UM | 1024 | strongly positive |
| 03-153 | UM | 256 | positive |
| 03-160 | UM | 1024 | strongly positive |

- **IgG anti-VSA detection and quantification by flow cytometry**

| sample | clinical group | result (RFU) |
|---|---|---|
| 01-004 | CM | 69750,055 |
| 01-006 | CM | 93936,055 |
| 01-013 | CM | 153647,71 |
| 01-020 | CM | 54280,015 |
| 02-002 | CM | 66634,795 |
| 02-003 | CM | 30863,04 |
| 02-011 | CM | 77202,57 |
| 02-013 | CM | 26680,735 |
| 02-016 | CM | 20623,13 |
| 02-017 | CM | 42574,78 |
| 02-018 | CM | 34028,605 |
| 02-025 | CM | 18176,735 |
| 02-027 | CM | 28147,13 |
| 02-030 | CM | 32834,095 |
| 02-034 | CM | 57273,3 |
| 03-022 | UM | 107062,375 |
| 03-025 | UM | 28354,09 |
| 03-031 | UM | 56882,875 |
| 03-037 | UM | 53430,445 |
| 03-059 | UM | 17200,115 |
| 03-060 | UM | 19994,2 |
| 03-075 | UM | 49401,25 |
| 03-110 | UM | 50234,22 |
| 03-113 | UM | 24078,77 |
| 03-137 | UM | 19633,59 |
| 03-140 | UM | 37966,305 |
| 03-141 | UM | 28721,45 |
| 03-145 | UM | 28208,495 |
| 03-153 | UM | 18823,67 |
| 03-160 | UM | 29646,6 |

#### Supplemental data 4: Parasites developmental blood stages

- **CM samples: microscopy**

| **Stage**<br><br>**(hpi)** | **Early ring**<br><br>**(0-16)** | **Late ring**<br>**+ Early trophozoite**<br><br>**(16-30)** | **Late**<br>**Trophozoite**<br><br>**(30-38)** | **Schizont**<br><br>**38-48** | **Gametocyte** |
|---|---|---|---|---|---|
| **01-004** | Not readable | | | | |
| **01-006** | Not readable | | | | |
| **01-013** | Not readable | | | | |
| **01-020** | 0.91 | 0.09 | 0 | 0 | 0 |
| **02-002** | 1 | 0 | 0 | 0 | 0 |
| **02-003** | 0.93 | 0.07 | 0 | 0 | 0 |
| **02-011** | 1 | 0 | 0 | 0 | 0 |
| **02-013** | 0.93 | 0.07 | 0 | 0 | 0 |
| **02-016** | Not readable | | | | |
| **02-017** | 1 | 0 | 0 | 0 | 0 |
| **02-018** | 0.94 | 0.06 | 0 | 0 | 0 |
| **02-025** | 0.96 | 0.04 | 0 | 0 | 0 |
| **02-027** | 0.98 | 0.02 | 0 | 0 | 0 |
| **02-030** | 0.55 | 0.45 | 0 | 0 | 0 |
| **02-034** | 1 | 0 | 0 | 0 | 0 |

- **CM samples: computed**

| **Stage**<br><br>**(hpi)** | **Ring**<br><br>**(8)** | **Early trophozoite**<br><br>**(19)** | **Late trophozoite**<br><br>**(30)** | **Schizont**<br><br>**(42)** | **Gametocyte** |
|---|---|---|---|---|---|
| **01-004** | 0.63 | 0.37 | 0 | 0 | 0 |
| **01-006** | 0.91 | 0.09 | 0 | 0 | 0 |
| **01-013** | 0.94 | 0.06 | 0 | 0 | 0 |
| **01-020** | 1 | 0 | 0 | 0 | 0 |
| **02-002** | 1 | 0 | 0 | 0 | 0 |
| **02-003** | 1 | 0 | 0 | 0 | 0 |
| **02-011** | 1 | 0 | 0 | 0 | 0 |
| **02-013** | 1 | 0 | 0 | 0 | 0 |
| **02-016** | 1 | 0 | 0 | 0 | 0 |
| **02-017** | 1 | 0 | 0 | 0 | 0 |
| **02-018** | 1 | 0 | 0 | 0 | 0 |
| **02-025** | 1 | 0 | 0 | 0 | 0 |
| **02-027** | 0.86 | 0.14 | 0 | 0 | 0 |
| **02-030** | 0.56 | 0.41 | 0.03 | 0 | 0 |
| **02-034** | 1 | 0 | 0 | 0 | 0 |

- **UM samples: microscopy**

| Stage (hpi) | Early ring (0-16) | Late ring + Early trophozoite (16-30) | Late Trophozoite (30-38) | Schizont 38-48 | Gametocyte |
|---|---|---|---|---|---|
| **03-022** | 0.99 | 0.01 | 0 | 0 | 0 |
| **03-025** | - | 1 | 0 | 0 | 0 |
| **03-031** | 0.01 | 0.99 | 0 | 0 | 0 |
| **03-037** | 0.21 | 0.79 | 0 | 0 | 0 |
| **03-059** | 0.01 | 0.99 | 0 | 0 | 0 |
| **03-060** | 0.44 | 0.56 | 0 | 0 | 0 |
| **03-075** | 0.06 | 0.94 | 0 | 0 | 0 |
| **03-110** | 0.86 | 0.14 | 0 | 0 | 0 |
| **03-113** | Not readable | | | | |
| **03-137** | 0.03 | 0.97 | 0 | 0 | 0 |
| **03-140** | - | 1 | 0 | 0 | 0 |
| **03-141** | 0.06 | 0.94 | 0 | 0 | 0 |
| **03-145** | 0.14 | 0.86 | 0 | 0 | 0 |
| **03-153** | 0.29 | 0.71 | 0 | 0 | 0 |
| **03-160** | 0.79 | 0.21 | 0 | 0 | 0 |

- **UM samples: computed**

| Stage (hpi) | Ring (8) | Early trophozoite (19) | Late trophozoite (30) | Schizont (42) | Gametocyte |
|---|---|---|---|---|---|
| **03-022** | 1 | 0 | 0 | 0 | 0 |
| **03-025** | 0.24 | 0.73 | 0 | 0 | 0.03 |
| **03-031** | 0.11 | 0.77 | 0.02 | 0 | 0.1 |
| **03-037** | 0.37 | 0.58 | 0 | 0 | 0.05 |
| **03-059** | 0.10 | 0.86 | 0 | 0 | 0.04 |
| **03-060** | 0.77 | 0.23 | 0 | 0 | 0 |
| **03-075** | 0.13 | 0.83 | 0 | 0 | 0.04 |
| **03-110** | 0.71 | 0.29 | 0 | 0 | 0 |
| **03-113** | 1 | 0 | 0 | 0 | 0 |
| **03-137** | 0.58 | 0.42 | 0 | 0 | 0 |
| **03-140** | 0.59 | 0.41 | 0 | 0 | 0 |
| **03-141** | 0.59 | 0.41 | 0 | 0 | 0 |
| **03-145** | 0.63 | 0.37 | 0 | 0 | 0 |
| **03-153** | 0.71 | 0.29 | 0 | 0 | 0 |
| **03-160** | 0.82 | 0.18 | 0 | 0 | 0 |

- **CM additional samples: microscopy**

| **Stage**<br><br>**(hpi)** | **Early ring**<br><br>**(0-16)** | **Late ring + Early trophozoite**<br><br>**(16-30)** | **Late Trophozoite**<br><br>**(30-38)** | **Schizont**<br><br>**38-48** | **Gametocyte** |
|---|---|---|---|---|---|
| **01-018** | 0.44 | 0.32 | 0.24 | 0 | 0 |
| **01-023** | 1 | 0 | 0 | 0 | 0 |
| **01-026** | 0.89 | 0.11 | 0 | 0 | 0 |
| **01-033** | 0.75 | 0.23 | 0.02 | 0 | 0 |
| **01-034** | 0.5 | 0.5 | 0 | 0 | 0 |
| **02-005** | 1 | 0 | 0 | 0 | 0 |
| **02-020** | 1 | 0 | 0 | 0 | 0 |
| **02-021** | 0.99 | 0 | 0 | 0.01 | 0 |
| **02-023** | 0.95 | 0.05 | 0 | 0 | 0 |
| **02-028** | 0.99 | 0.01 | 0 | 0 | 0 |
| **02-038** | 1 | 0 | 0 | 0 | 0 |
| **02-039** | 0.98 | 0.02 | 0 | 0 | 0 |
| **02-043** | 1 | 0 | 0 | 0 | 0 |
| **02-044** | 1 | 0 | 0 | 0 | 0 |
| **02-049** | 1 | 0 | 0 | 0 | 0 |

- **UM additional samples: microscopy**

| **Stage**<br><br>**(hpi)** | **Early ring**<br><br>**(0-16)** | **Late ring + Early trophozoite**<br><br>**(16-30)** | **Late Trophozoite**<br><br>**(30-38)** | **Schizont**<br><br>**38-48** | **Gametocyte** |
|---|---|---|---|---|---|
| **03-039** | 1 | 0 | 0 | 0 | 0 |
| **03-047** | 0.19 | 0.81 | 0 | 0 | 0 |
| **03-063** | 0.24 | 0.76 | 0 | 0 | 0 |
| **03-093** | 1 | 0 | 0 | 0 | 0 |
| **03-095** | 1 | 0 | 0 | 0 | 0 |
| **03-100** | 0.89 | 0.11 | 0 | 0 | 0 |
| **03-101** | 0.56 | 0.44 | 0 | 0 | 0 |
| **03-105** | 0.12 | 0.88 | 0 | 0 | 0 |
| **03-116** | 0.27 | 0.73 | 0 | 0 | 0 |
| **03-122** | 0.77 | 0.23 | 0 | 0 | 0 |
| **03-125** | 0.61 | 0.30 | 0 | 0 | 0.09 |
| **03-132** | 0.47 | 0.53 | 0 | 0 | 0 |
| **03-138** | 0.99 | 0.01 | 0 | 0 | 0 |
| **03-144** | 0.14 | 0.86 | 0 | 0 | 0 |
| **03-154** | 1 | 0 | 0 | 0 | 0 |

### Supplemental data 5: RT-qPCR validations

- **Sequenced group**

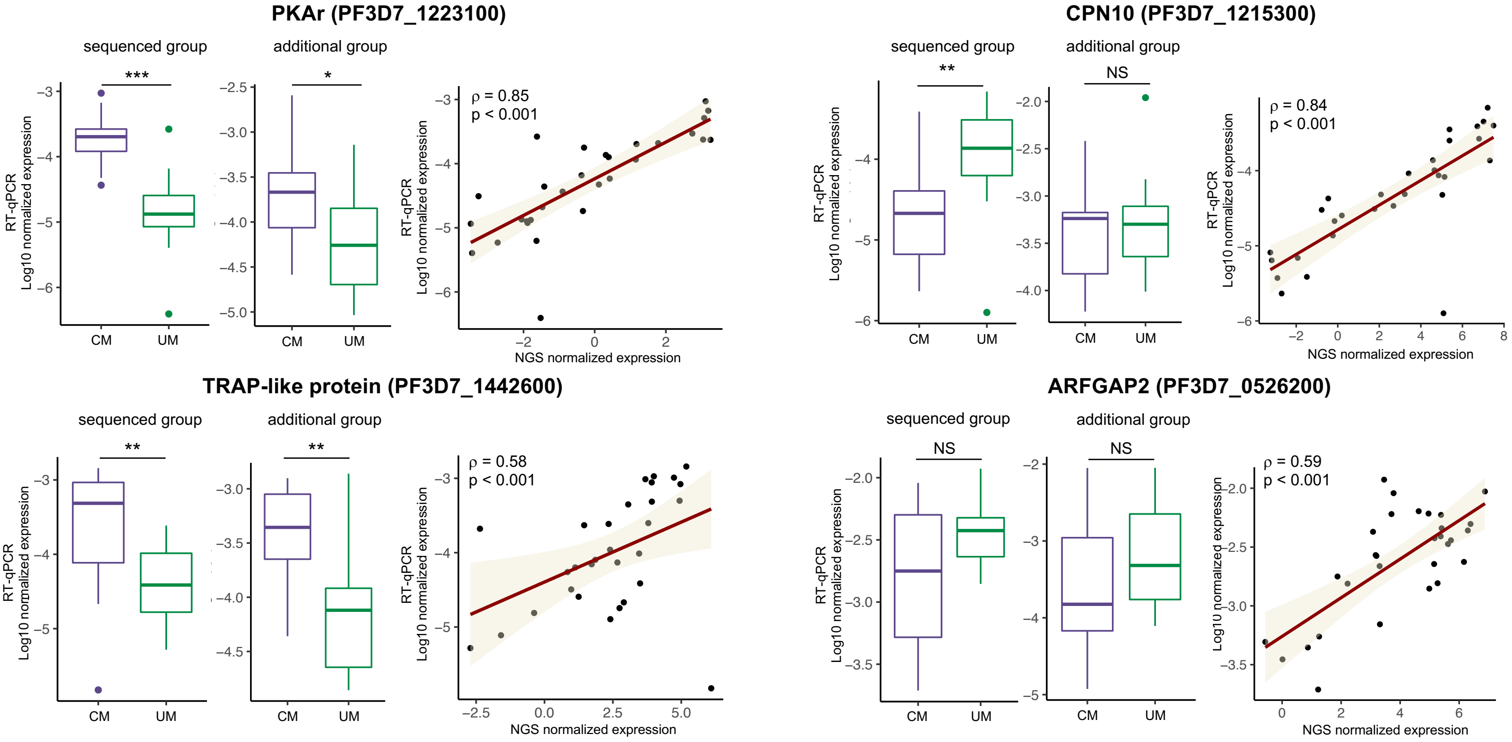

| Gene | **CM**<br>Median [10th-90th percentile] | **UM**<br>Median [10th-90th percentile] | **p-value** |
|---|---|---|---|
| PHISTa - PF3D7_1372000 | $1.1*10^{-4}$ [$3.6*10^{-5}$-$5.9*10^{-4}$] | $3.0*10^{-4}$ [$2.0*10^{-4}$-$6.1*10^{-4}$] | < 0.001 |
| PKAr - PF3D7_1223100 | $2.0*10^{-4}$ [$5.2*10^{-5}$-$6.1*10^{-4}$] | $1.3*10^{-5}$ [$4.8*10^{-6}$-$5.7*10^{-5}$] | < 0.001 |
| CLAG9 - PF3D7_0935800 | $5.5*10^{-4}$ [$1.6*10^{-4}$-$1.5*10^{-3}$] | $9.3*10^{-5}$ [$3.1*10^{-5}$-$1.1*10^{-3}$] | 0.02 |
| TRAP-like protein - PF3D7_1442600 | $4.9*10^{-4}$ [$3.8*10^{-5}$-$1.0*10^{-3}$] | $3.9*10^{-5}$ [$9.8*10^{-6}$-$2.2*10^{-4}$] | 0.002 |
| HSP60 - PF3D7_1015600 | $1.6*10^{-3}$ [$5.3*10^{-4}$-$2.1*10^{-2}$] | $2.7*10^{-4}$ [$9.6*10^{-5}$-$6.3*10^{-4}$] | < 0.001 |
| CPN10 - PF3D7_1215300 | $2.1*10^{-5}$ [$3.8*10^{-6}$-$7.5*10^{-5}$] | $1.4*10^{-4}$ [$3.5*10^{-5}$-$4.3*10^{-4}$] | 0.002 |
| Ppase - PF3D7_0316300 | $3.9*10^{-5}$ [$1.4*10^{-5}$-$2.8*10^{-4}$] | $9.6*10^{-4}$ [$1.8*10^{-4}$-$1.8*10^{-3}$] | < 0.001 |
| ARFGAP2 - PF3D7_0526200 | $1.8*10^{-3}$ [$3.9*10^{-4}$-$6.1*10^{-3}$] | $3.8*10^{-3}$ [$1.8*10^{-3}$-$8.2*10^{-3}$] | 0.09 |

- **Additional group**

| **Gene** | **CM** Median [10th-90th percentile] | **UM** Median [10th-90th percentile] | **p-value** |
|---|---|---|---|
| PHISTa - PF3D7_1372000 | $9.2*10^{-4}$ [$1.9*10^{-4}$ - $4.4*10^{-2}$] | $1.1*10^{-4}$ [$6.6*10^{-5}$ - $2.5*10^{-3}$] | 0.004 |
| PKAr - PF3D7_1223100 | $2.1*10^{-4}$ [$3.9*10^{-5}$ -$1.1*10^{-3}$] | $5.5*10^{-5}$ [$1.3*10^{-5}$ - $4.9*10^{-4}$] | 0.03 |
| CLAG9 - PF3D7_0935800 | $3.9*10^{-4}$ [$3.0*10^{-5}$ - $1.7*10^{-3}$] | $1.0*10^{-4}$ [$2.1*10^{-5}$ - $5.4*10^{-4}$] | 0.02 |
| TRAP-like protein - PF3D7_1442600 | $4.4*10^{-4}$ [$9.0*10^{-5}$ - $1.1*10^{-3}$] | $7.6*10^{-5}$ [$1.8*10^{-5}$ - $3.3*10^{-4}$] | 0.001 |
| HSP60 - PF3D7_1015600 | $2.2*10^{-4}$ [$5.9*10^{-5}$ - $1.1*10^{-3}$] | $2.5*10^{-4}$ [$7.9*10^{-5}$ - $6.9*10^{-4}$] | 0.9 |
| CPN10 - PF3D7_1215300 | $5.8*10^{-4}$ [$7.4*10^{-5}$ - $1.7*10^{-3}$] | $5.0*10^{-4}$ [$1.6*10^{-4}$ - $1.3*10^{-3}$] | 0.6 |
| Ppase - PF3D7_0316300 | $1.3*10^{-4}$ [$3.6*10^{-5}$ - $2.2*10^{-3}$] | $4.3*10^{-4}$ [$1.1*10^{-4}$ - $3.8*10^{-3}$] | 0.2 |
| ARFGAP2 - PF3D7_0526200 | $1.5*10^{-4}$ [$4.0*10^{-5}$ - $4.0*10^{-3}$] | $4.8*10^{-4}$ [$1.2*10^{-4}$ - $4.4*10^{-3}$] | 0.2 |

#### Supplemental data 6: GSEA – GO biological process and KEGG pathways

- **GO biological process**

| **GO:0051276** | **chromosome organization** |
|---|---|
| PF3D7_0610400 | histone H3 |
| PF3D7_0617800 | histone H2A |
| PF3D7_0617900 | histone H3 variant |
| PF3D7_0509100 | structural maintenance of chromosomes protein 4, putative |
| PF3D7_0414000 | structural maintenance of chromosomes protein 3 |
| PF3D7_1318400 | structural maintenance of chromosomes protein 2, putative |
| **GO:1902600** | **proton transmembrane transport** |
| PF3D7_1456800 | V-type H(+)-translocating pyrophosphatase, putative |
| PF3D7_1323200 | V-type proton ATPase subunit G, putative |
| PF3D7_0217100 | ATP synthase subunit alpha, mitochondrial |
| PF3D7_1235200 | V-type K+-independent H+-translocating inorganic pyrophosphatase |
| PF3D7_1235700 | ATP synthase subunit beta, mitochondrial |
| **GO:0006412** | **translation** |
| PF3D7_1223900 | 50S ribosomal protein L24, putative |
| PF3D7_1442800 | conserved Plasmodium protein, unknown function |
| PF3D7_1144000 | 40S ribosomal protein S21 |
| PF3D7_1414300 | 60S ribosomal protein L10, putative |
| PF3D7_0611700 | 60S ribosomal protein L39 |
| PF3D7_1144300 | 60S ribosomal protein L41 |
| PF3D7_0710600 | 60S ribosomal protein L34 |
| PF3D7_1351400 | 60S ribosomal protein L17, putative |
| PF3D7_1461300 | 40S ribosomal protein S28e, putative |
| PF3D7_0317600 | 40S ribosomal protein S11, putative |
| PF3D7_1317800 | 40S ribosomal protein S19 |
| PF3D7_0316800 | 40S ribosomal protein S15A, putative |
| PF3D7_1308300 | 40S ribosomal protein S27 |
| PF3D7_1408600 | 40S ribosomal protein S8e, putative |
| PF3D7_1431700 | 60S ribosomal protein L14, putative |
| PF3D7_0415900 | 60S ribosomal protein L15, putative |
| PF3D7_0212200 | ribosomal protein L12, mitochondrial, putative |
| PF3D7_0706400 | 60S ribosomal protein L37 |
| PF3D7_0312800 | 60S ribosomal protein L26, putative |
| PF3D7_0304400 | 60S ribosomal protein L44 |
| PF3D7_1124900 | 60S ribosomal protein L35, putative |
| PF3D7_1331800 | 60S ribosomal protein L23, putative |
| PF3D7_0814000 | 60S ribosomal protein L13-2, putative |
| PF3D7_1242700 | 40S ribosomal protein S17, putative |
| PF3D7_0422400 | 40S ribosomal protein S19 |
| PF3D7_0306900 | 40S ribosomal protein S23, putative |
| PF3D7_1003500 | 40S ribosomal protein S20e, putative |
| PF3D7_1105400 | 40S ribosomal protein S4, putative |
| PF3D7_0307200 | 60S ribosomal protein L7, putative |

| | |
|---|---|
| PF3D7_0317100 | 6-cysteine protein |
| PF3D7_0610000 | ribosomal protein L19, mitochondrial, putative |
| PF3D7_0822000 | ribosomal protein L4, mitochondrial, putative |
| **GO:0006270** | **DNA replication initiation** |
| PF3D7_1425900 | conserved Plasmodium protein, unknown function |
| PF3D7_1355100 | DNA replication licensing factor MCM6 |
| PF3D7_1211700 | DNA replication licensing factor MCM5, putative |
| PF3D7_1317100 | DNA replication licensing factor MCM4 |
| PF3D7_1417800 | DNA replication licensing factor MCM2 |
| **GO:0006260** | **DNA replication** |
| PF3D7_1425900 | conserved Plasmodium protein, unknown function |
| PF3D7_1355100 | DNA replication licensing factor MCM6 |
| PF3D7_1463200 | replication factor C subunit 3, putative |
| PF3D7_1334100 | conserved protein, unknown function |
| PF3D7_1111100 | replication factor C subunit 5, putative |
| PF3D7_0411900 | DNA polymerase alpha catalytic subunit A |
| PF3D7_1211700 | DNA replication licensing factor MCM5, putative |
| PF3D7_1317100 | DNA replication licensing factor MCM4 |
| PF3D7_1015800 | ribonucleoside-diphosphate reductase small chain, putative |
| **GO:0006414** | **translational elongation** |
| PF3D7_1442800 | conserved Plasmodium protein, unknown function |
| PF3D7_0309600 | 60S acidic ribosomal protein P2 |
| PF3D7_1235400 | tetQ family GTPase, putative |
| PF3D7_0110100 | selenocysteine-specific elongation factor, putative |
| PF3D7_1330600 | elongation factor Tu, putative |
| **GO:0007018** | **microtubule-based movement** |
| PF3D7_1020300 | cytoplasmic dynein intermediate chain, putative |
| PF3D7_1122900 | dynein heavy chain, putative |
| PF3D7_1211000 | kinesin-X3, putative |
| PF3D7_0319400 | kinesin-8X |
| PF3D7_1426300 | dynein intermediate chain, putative |
| PF3D7_0729900 | dynein heavy chain, putative |
| **GO:0006352** | **DNA-templated transcription, initiation** |
| PF3D7_1404000 | DNA-directed RNA polymerase II subunit RPB4, putative |
| PF3D7_0621000 | conserved Plasmodium protein, unknown function |
| PF3D7_0506200 | TATA-box-binding protein |
| **GO:0006333** | **chromatin assembly or disassembly** |
| PF3D7_0610400 | histone H3 |
| PF3D7_0320900 | histone H2A.Z |
| PF3D7_1220900 | heterochromatin protein 1 |
| **GO:0044409** | **entry into host** |
| PF3D7_0404400 | 6-cysteine protein P36 |
| PF3D7_0309600 | 60S acidic ribosomal protein P2 |
| PF3D7_1133400 | apical membrane antigen 1 |
| PF3D7_1136700 | armadillo-interacting protein AIP |

| | |
|---|---|
| PF3D7_1458000 | cysteine proteinase falcipain 1 |
| PF3D7_0731500 | erythrocyte binding antigen-175 |
| PF3D7_0102500 | erythrocyte binding antigen-181 |
| PF3D7_1147800 | membrane associated erythrocyte binding-like protein |
| PF3D7_1430200 | plasmepsin IX |
| PF3D7_0808200 | plasmepsin X |
| PF3D7_1335400 | reticulocyte binding protein 2 homologue a |
| PF3D7_0424200 | reticulocyte binding protein homologue 4 |
| PF3D7_1116000 | rhoptry neck protein 4 |
| PF3D7_0501600 | rhoptry-associated protein 2 |
| **GO:0048193** | **Golgi vesicle transport** |
| PF3D7_1437800 | trafficking protein particle complex subunit 5, putative |
| PF3D7_0418500 | trafficking protein particle complex subunit 3, putative |
| **GO:0019752** | **carboxylic acid metabolic process** |
| PF3D7_1325200 | lactate dehydrogenase, putative |
| PF3D7_0618500 | malate dehydrogenase |
| **GO:0009657** | **plastid organization** |
| PF3D7_1239700 | ATP-dependent zinc metalloprotease FTSH 1 |
| PF3D7_1408700 | conserved protein, unknown function |
| PF3D7_0913500 | protease, putative |
| PF3D7_1417800 | DNA replication licensing factor MCM2 |

- **KEGG pathways**

| **ec00010** | **Glycolysis / Gluconeogenesis** |
|---|---|
| PF3D7_0303700 | lipoamide acyltransferase component of branched-chain alpha-keto acid dehydrogenase complex |
| PF3D7_1342800 | malate dehydrogenase |
| PF3D7_0922500 | ATP-dependent 6-phosphofructokinase |
| PF3D7_0915400 | phosphoglycerate kinase |
| PF3D7_1325200 | lactate dehydrogenase, putative |
| PF3D7_1446400 | phosphoenolpyruvate carboxykinase |
| PF3D7_0618500 | pyruvate dehydrogenase E1 component subunit beta |
| **ec00750** | **Vitamin B6 metabolism** |
| PF3D7_0616000 | pyridoxal kinase |
| PF3D7_1459700 | pyridoxal 5'-phosphate synthase, putative |
| **ec00620** | **Pyruvate metabolism** |
| PF3D7_0303700 | lipoamide acyltransferase component of branched-chain alpha-keto acid dehydrogenase complex |
| PF3D7_0618500 | malate dehydrogenase |
| PF3D7_1325200 | lactate dehydrogenase, putative |
| PF3D7_1342800 | phosphoenolpyruvate carboxykinase |
| PF3D7_1446400 | pyruvate dehydrogenase E1 component subunit beta |
| PF3D7_1450900 | acetyl-CoA acetyltransferase, putative |

#### Supplemental data 7: MCode clusters from PPI network

| Cluster 1 | |
|---|---|
| PF3D7_0304400 | 60S ribosomal protein L44 |
| PF3D7_0306900 | 40S ribosomal protein S23, putative |
| PF3D7_0307200 | 60S ribosomal protein L7, putative |
| PF3D7_0309600 | 60S acidic ribosomal protein P2 |
| PF3D7_0312800 | 60S ribosomal protein L26, putative |
| PF3D7_0316800 | 40S ribosomal protein S15A, putative |
| PF3D7_0317600 | 40S ribosomal protein S11, putative |
| PF3D7_0415900 | 60S ribosomal protein L15, putative |
| PF3D7_0422400 | 40S ribosomal protein S19 |
| PF3D7_0611700 | 60S ribosomal protein L39 |
| PF3D7_0706400 | 60S ribosomal protein L37 |
| PF3D7_0710600 | 60S ribosomal protein L34 |
| PF3D7_0814000 | 60S ribosomal protein L13-2, putative |
| PF3D7_1003500 | 40S ribosomal protein S20e, putative |
| PF3D7_1105400 | 40S ribosomal protein S4, putative |
| PF3D7_1124900 | 60S ribosomal protein L35, putative |
| PF3D7_1144000 | 40S ribosomal protein S21 |
| PF3D7_1242700 | 40S ribosomal protein S17, putative |
| PF3D7_1308300 | 40S ribosomal protein S27 |
| PF3D7_1317800 | 40S ribosomal protein S19 |
| PF3D7_1331800 | 60S ribosomal protein L23, putative |
| PF3D7_1351400 | 60S ribosomal protein L17, putative |
| PF3D7_1408600 | 40S ribosomal protein S8e, putative |
| PF3D7_1414300 | 60S ribosomal protein L10, putative |
| PF3D7_1431700 | 60S ribosomal protein L14, putative |
| PF3D7_1461300 | 40S ribosomal protein S28e, putative |
| **Cluster 2** | |
| PF3D7_0508000 | 6-cysteine protein |
| PF3D7_1133400 | apical membrane antigen 1 |
| PF3D7_0611600 | basal complex transmembrane protein 1 |
| PF3D7_1223100 | cAMP-dependent protein kinase regulatory subunit |
| PF3D7_0621100 | conserved Plasmodium protein, unknown function |
| PF3D7_1136200 | conserved Plasmodium protein, unknown function |
| PF3D7_1206300 | conserved Plasmodium protein, unknown function |
| PF3D7_1435600 | conserved Plasmodium protein, unknown function |
| PF3D7_0723300 | conserved protein, unknown function |
| PF3D7_0817600 | conserved protein, unknown function |
| PF3D7_1320700 | conserved protein, unknown function |
| PF3D7_0302500 | cytoadherence linked asexual protein 3.1 |
| PF3D7_0302200 | cytoadherence linked asexual protein 3.2 |
| PF3D7_0831600 | cytoadherence linked asexual protein 8 |
| PF3D7_1035300 | glutamate-rich protein GLURP |
| PF3D7_1345600 | inner membrane complex protein |

| | |
|---|---|
| PF3D7_0613900 | myosin E, putative |
| PF3D7_0822900 | PhIL1-interacting candidate PIC2 |
| PF3D7_0508900 | protein AAP6 |
| PF3D7_1321100 | protein kinase domain-containing protein, putative |
| PF3D7_0932100 | protein MAM3, putative |
| PF3D7_0506900 | rhomboid protease ROM4 |
| PF3D7_1463900 | rhoptry neck protein 11, putative |
| PF3D7_1452000 | rhoptry neck protein 2 |
| PF3D7_0817700 | rhoptry neck protein 5 |
| PF3D7_1410400 | rhoptry-associated protein 1 |
| PF3D7_0501600 | rhoptry-associated protein 2 |
| PF3D7_0207800 | serine repeat antigen 3 |
| PF3D7_1356800 | serine/threonine protein kinase ARK3, putative |
| PF3D7_0704500 | serine/threonine protein kinase, putative |
| PF3D7_0508100 | SET domain protein, putative |
| PF3D7_1235200 | V-type K+-independent H+-translocating inorganic pyrophosphatase |
| **Cluster 3** | |
| PF3D7_0102500 | erythrocyte binding antigen-181 |
| PF3D7_0203100 | protein kinase, putative |
| PF3D7_0808200 | plasmepsin X |
| PF3D7_1125800 | kelch domain-containing protein, putative |
| PF3D7_1252400 | reticulocyte binding protein homologue 3, pseudogene |
| PF3D7_1332200 | conserved protein, unknown function |
| PF3D7_1335400 | reticulocyte binding protein 2 homologue a |
| **Cluster 4** | |
| PF3D7_0316300 | inorganic pyrophosphatase |
| PF3D7_0319400 | kinesin-8X |
| PF3D7_0516800 | AP2 domain transcription factor AP2-O2, putative |
| PF3D7_0707200 | conserved Plasmodium protein, unknown function |
| PF3D7_0717600 | inner membrane complex protein, putative |
| PF3D7_0815500 | conserved Plasmodium protein, unknown function |
| PF3D7_1133200 | conserved Plasmodium protein, unknown function |
| PF3D7_1211700 | DNA replication licensing factor MCM5, putative |
| PF3D7_1239200 | AP2 domain transcription factor, putative |
| PF3D7_1317100 | DNA replication licensing factor MCM4 |
| PF3D7_1355100 | DNA replication licensing factor MCM6 |
| PF3D7_1417800 | DNA replication licensing factor MCM2 |

#### Supplemental data 8: *var* genes expression measurement by RT-qPCR

- **Sequenced group**

| **Primers set** | **CM**<br>Median [10th-90th percentile] | **UM**<br>Median [10th-90th percentile] | **p-value** |
|---|---|---|---|
| ATS-1 | 0.004 [0.002-0.008] | 0.001 [0.0006-0.003] | <0.001 |
| ATS-2 | 0.02 [0.01-0.05] | 0.01 [0.008-0.02] | <0.001 |
| DBLα | 0.1 [0.02-0.9] | 0.05 [0.02-0.3] | 0.1 |

- **Additional group**

| **Primers set** | **CM**<br>Median [10th-90th percentile] | **UM**<br>Median (10th-90th percentile) | **p-value** |
|---|---|---|---|
| ATS-1 | 0.002 [0.0006-0.01] | 0.001 [0.0003-0.002] | 0.08 |
| ATS-2 | 0.08 [0.04-0.5] | 0.04 [0.02-0.3] | 0.03 |
| DBLα | 0.07 [0.02-0.2] | 0.06 [0.02-0.2] | 0.7 |

#### Supplemental data 9: *var* ATS-2 primers

| | Primers set | Vardom statistics | VarDB statistics |
|---|---|---|---|
| **Forward** | cccatccacaaccaactgga<br>ccaattatgaaccaattaga<br>cctatactcaatcaaataaa<br>cctataaccaatcaaataaa | 47%<br>(106/226) | 13%<br>(196/1517) |
| **Reverse** | agatagacatagaratatgtgyga<br>agatagacatagagatatgtgcga<br>agatagacatagagatatgtgtga<br>agatagacatagaaatatgtgtga<br>agatagacatagaaatatgtgcga | 68%<br>(154/226) | 46%<br>(699/1517) |
